# Loss of ABCA4 from photoreceptor discs triggers changes in glial cell homeostasis

**DOI:** 10.64898/2026.04.13.718110

**Authors:** Rossella Valenzano, Andrew McDonald, Carmen Gallego, Charlotte A. Andriessen, Ioannis Moustakas, Aat A. Mulder, Harald M.M. Mikkers, Roman I. Koning, Hailiang Mei, Jan Wijnholds

**Author notes:** Co-first author (Rossella Valenzano, Andrew McDonald, Carmen Gallego). VectorY Therapeutics, Science Park 408, 1098 XH Amsterdam, The Netherlands. **Author Contributions:** Rossella Valenzano: Conception and design, collection and/or assembly of data, data analysis and interpretation, manuscript writing, final approval of manuscript Andrew McDonald: Conception and design, collection and/or assembly of data, data analysis and interpretation, final approval of manuscript Carmen Gallego: Conception and design, collection and/or assembly of data, data analysis and interpretation, final approval of manuscript Charlotte A. Andriessen: Collection and/or assembly of data, data analysis and interpretation, final approval of manuscript Ioannis Moustakas: Collection and/or assembly of data, data analysis and interpretation, final approval of manuscript Aat A. Mulder: Collection and/or assembly of data, data analysis and interpretation, final approval of manuscript Harald M.M. Mikkers: Conception and design, final approval of manuscript Roman I. Koning: Collection and/or assembly of data, data analysis and interpretation, final approval of manuscript Hailiang Mei: Collection and/or assembly of data, data analysis and interpretation, final approval of manuscript Jan Wijnholds: Conception and design, funding acquisition, administrative support, final approval of manuscript.

## Abstract

Loss-of-function mutations in the *ABCA4* gene cause Stargardt disease (STGD1), the most common inherited macular dystrophy leading to progressive central vision loss. Here, we generated hiPSC-derived retinal organoids harboring a premature stop codon in exon-24 of *ABCA4* to evaluate the impact of this mutation on mRNA and protein levels in a human model. Immunofluorescence analysis revealed the absence of ABCA4 protein in the mutant photoreceptor outer segment discs, while single-cell RNA sequencing detected no major transcriptional alterations in rods and cones. Unexpectedly, differential gene expression and pathway enrichment analyses of Müller glial cells (MGCs) and astrocytes highlighted disruption of neuronal development, microenvironment of glial cells, intercellular communication, and programmed cell death pathways. These findings suggest that *ABCA4* might play a role in maintaining the retinal microenvironment homeostasis, and that the early transcriptomic response of MGCs and astrocytes preceding photoreceptor degeneration could contribute to Stargardt disease development.

**Significance Statement:** Human induced pluripotent stem cell (hiPSC)-derived retinal organoids provide a powerful platform to investigate inherited retinal diseases. In this study, we generated *ABCA4*-mutant hiPSC lines and differentiated them into retinal organoids to model Stargardt disease. Despite complete loss of ABCA4 protein from the photoreceptor outer segment discs, rods and cones exhibited minimal transcriptional alterations. In contrast, the *ABCA4* variant triggered changes in the glial cell homeostasis, suggesting that Müller glial cells and astrocytes might exhibit an early response to photoreceptor dysfunction in the absence of ABCA4.

Human induced pluripotent stem cells (hiPSCs) were engineered to generate *ABCA4*-mutant cell lines, later differentiated into retinal organoids as a model of Stargardt disease. The organoids showed loss of ABCA4 from the outer segment discs of rod and cone photoreceptors, while the mutant Müller glial cells and astrocytes exhibited transcriptional changes in pathways involved in neuronal development, microenvironment, and programmed cell death.

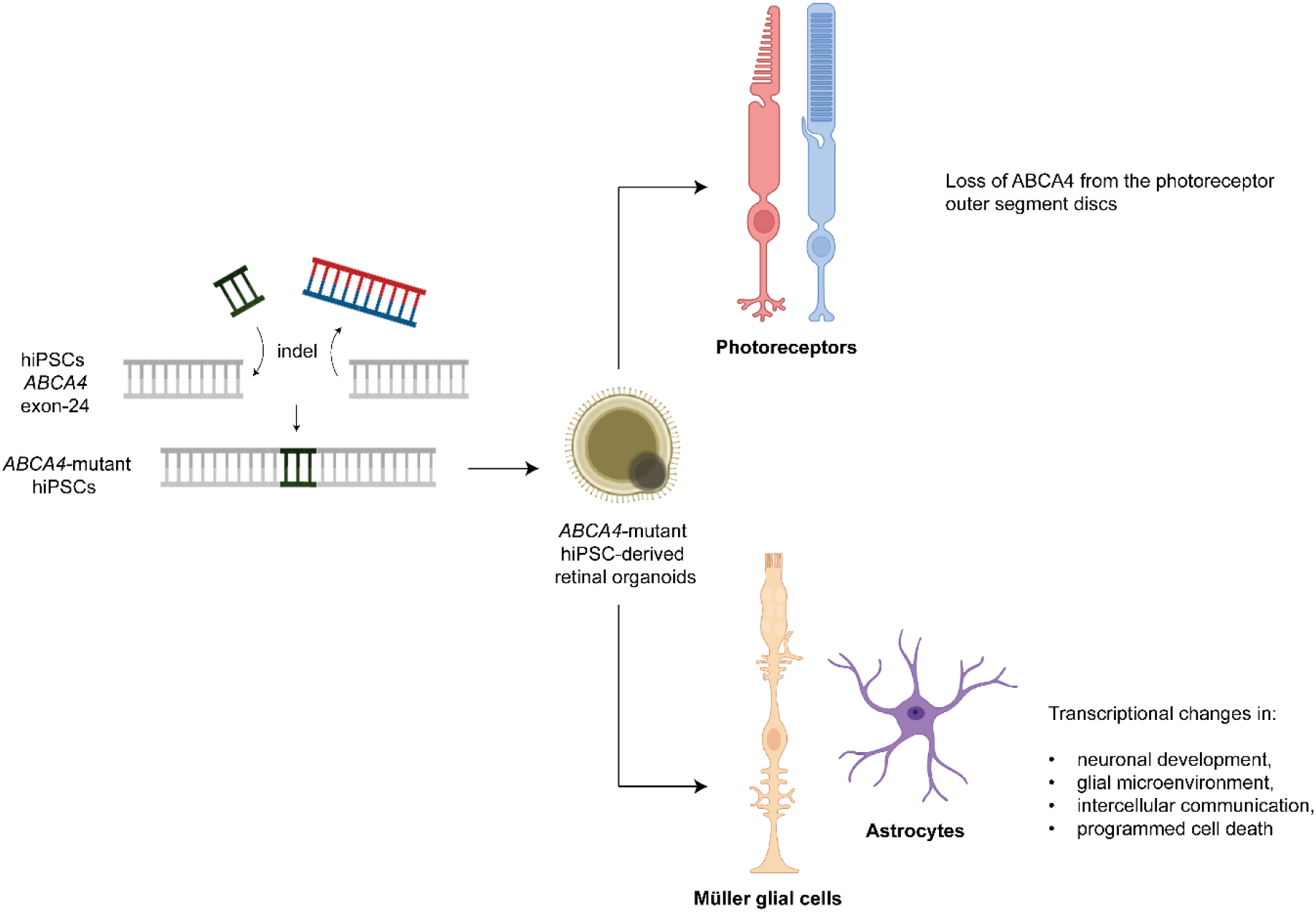

## Introduction

The ATP-binding cassette, sub-family A, member 4 protein, also known as ABCA4, plays a critical role in the recycling of metabolites in the visual cycle. ABCA4 is predominantly localized to the membranous discs of photoreceptor outer segments, where it acts as a flippase, transporting N-retinylidene phosphatidylethanolamine (N-Ret-PE) from the luminal to the cytoplasmic leaflet of disc membranes, allowing the clearance of potentially toxic compounds from the photoreceptors [1-3]. Loss of function of ABCA4 protein leads to the accumulation of N-Ret-PE in the outer segment discs, where it is converted to phosphatidyl-pyridinium bisretinoid (A2PE). Following disc shedding and phagocytosis by the retinal pigment epithelium (RPE), A2PE is hydrolyzed to the toxic bisretinoid A2E [4, 5], whose accumulation contributes to RPE dysfunction and subsequent photoreceptor degeneration [6, 7].

*ABCA4* mutations are commonly associated with Stargardt disease (STGD1), an autosomal recessive retinal dystrophy causing vision loss in children and young adults. It has a reported prevalence of 1 in 7000, and a high frequency of variant carriers in the general population [8, 9]. Mutations in the *ABCA4* gene are also less frequently implicated in other retinal disorders falling outside the canonical classification of STGD1 [10]. *ABCA4* is located on the short arm of chromosome 1, where its 50 exons encode a large transmembrane protein of 2273 amino acids [11, 12]. *ABCA4* is known to have more than 1200 disease-associated variants, resulting in a broad range of heterogeneous phenotypes [5, 13]. Currently, there is no cure for *ABCA4*-related retinal dystrophies, and the high genotypic and phenotypic variability has posed a challenge for accurate disease modelling.

Mice are the most widely used *in vivo* model for STGD1, particularly *Abca4* knockout (*Abca4* ^-^/^-^) and disease-specific knock-in strains that recapitulate patient mutations [14-18]. These models have been instrumental in elucidating the role of ABCA4 transporter in the visual cycle, the accumulation of toxic bisretinoids such as A2E, and the subsequent formation of lipofuscin within the RPE. They have also served as key platforms to test therapeutic strategies [14, 16, 19]. However, mice lack a macula and possess a rod-dominant retina with a relatively low proportion of cone photoreceptors, limiting their ability to fully model the macular degeneration and cone dysfunction that characterize human STGD1. To address these anatomical and physiological differences, additional animal models have been developed. Although rats also lack a true macula, their retinal organization and larger eye size offer practical advantages for surgical manipulation and *in vivo* imaging in preclinical therapeutic studies [20, 21]. Zebrafish have emerged as a complementary model due to their cone-rich retina, which more closely mirrors the human retina composition [22-24]. Canine models represent a highly valuable large-animal system for STGD1 research. In fact, dogs present a cone-enriched region known as the area centralis, that is functionally analogous to the human macula [25]. Even pig models have been developed and used to test retinal gene therapy with dual adeno-associated viral (AAV) intein vectors [26], achieving better results than those obtained via other dual AAV-based strategies [27].

Recent developments in stem cell technology have allowed for the differentiation of human induced pluripotent stem cells (hiPSCs) into 3D retinal organoids that closely resemble human retinal architecture, with well organized, stratified tissue that contains all major retinal cell types [28, 29]. Retinal organoids have been used to model a number of inherited retinal dystrophies, offering a unique opportunity to study the underlying pathological mechanisms of retinal diseases in a human-derived system [30-33]. These retinal organoids contain both rod and cone photoreceptors with rudimentary outer-segments and disc-like structures, representing a valuable tool for the investigation of Stargardt disease and other *ABCA4*-associated disorders [34, 35]. This powerful system, paired with CRISPR/Cas9 technology, has facilitated the creation of precise knock-out models for the study of loss-of-function mutations. Retinal organoids with ablated proteins of interest offer an excellent platform to test potential gene therapies for the reintroduction of functional proteins, including viral-delivered gene augmentation or gene editing approaches [32, 36, 37].

In this study, we used CRISPR/Cas9 to introduce a premature stop codon in exon-24 of *ABCA4* in human iPSCs. The resulting *ABCA4* hiPSC-derived retinal organoids constitute a valuable disease model for future testing of homology-independent targeted integration (HITI)-based [38] *ABCA4* gene editing strategies. The mutant cell lines were differentiated into 3D retinal organoids and showed loss of ABCA4 protein from the photoreceptor discs. Single-cell RNA sequencing data revealed an unexpected response of Müller glial cells and astrocytes to the mutation, with transcriptional changes suggestive of altered regulation of apoptosis, cell-to-cell communication, and neuronal development pathways. These data provide new insights into the potential role of glial cells as therapeutic targets for the treatment of Stargardt disease.

## Materials and Methods

### hiPSC genome editing by CRISPR/Cas9

The gRNA targeting exon-24 of *ABCA4* was selected using the integrated DNA technology (IDT) Custom Alt-R™ CRISPR-Cas9 guide RNA online tool (https://eu.idtdna.com/). A 153bp single-stranded oligodeoxynucleotide (ssODN) HDR template (IDT, Madison, WI, USA) was designed to introduce an in-frame premature stop codon followed by a 10bp deletion. The SpCas9 Nuclease V3 protein high fidelity (IDT, Madison, WI, USA) was used to generate a ribonucleoprotein (RNP) complex. Genome editing was performed as previously described [39]. The parental LUMC0004iCTRL10 hiPSCs [40] were transfected with the Cas9-RNP complex and the ssODN using the Neon Transfection System (ThermoFisher Scientific, Waltham, MA, USA) with pulse conditions set to 1200 V/30 ms/1 pulse. Cells were then plated on Matrigel-coated plates (Corning, Glendale, AZ, USA) with mTeSR plus medium (STEMCELL Technologies, Cologne, Germany) supplemented with CloneR (STEMCELL Technologies, Cologne, Germany). After a recovery period, 1000 single cells were seeded on Matrigel-coated 10 cm dishes for subsequent selection of single colonies for genotyping. The DNA of clones was extracted using QuickExtract DNA extraction solution (Merck, Schiphol-Rijk, The Netherlands), and the region of interest was amplified by PCR. *ABCA4* subclones were confirmed by Sanger sequencing. Additional information on the hiPSC lines and the Materials used can be found in Tables S1-S3.

### Cell Culture and Retinal Organoid Differentiation

hiPSCs were cultured on Matrigel-coated 6-well plates in mTeSR plus and incubated at 37°C and 5% CO_2._ Cells were passaged following incubation with Gentle Cell Dissociation reagent (GCDR) (STEMCELL Technologies, Cologne, Germany) and mechanical scraping. Retinal organoid differentiation followed an established protocol [39]. In short, after three weeks of culture, hiPSCs were resuspended in mTeSR plus supplemented with 10 µM blebbistatin (Abcam, Waltham, MA, USA). One million cells were seeded per agarose micro-mold (Merck, Schiphol-Rijk, The Netherlands) in a medium gradually transitioned to NIM1. Embryoid bodies (EBs) were transferred to Matrigel-coated 6-well plates in NIM1. From DD10, 100 nM of SAG (Selleck Chemicals, Houston, TX, USA) was added, and from DD16 cultures were maintained in NIM2 with SAG. Between DD21 and DD28, neuroepithelial structures were manually isolated and cultured in agarose-coated 48-well plates in NIM2. From DD35 to DD44, organoids were cultured in RLM1 + 1 µM retinoic acid (Merck, Schiphol-Rijk, The Netherlands), followed by RLM1 supplemented with 1 µM retinoic acid and 10 µM DAPT (Selleck Chemicals, Houston, TX, USA) until DD55. From DD56 to DD85, organoids were maintained in RLM1 with retinoic acid, after which cultures were switched to RLM2 with 0.5 µM retinoic acid. From DD120 onward, organoids were maintained in RLM2 until collection. Additional information on the Materials used can be found in Table S3.

### Immunofluorescence analysis

Organoids were fixed with 4% paraformaldehyde in PBS for 20 minutes followed by a brief washing in PBS. Fixed organoids were cryopreserved in 15% sucrose PBS solution for 30 minutes at RT, followed by incubation in a 30% sucrose PBS solution for at least 1 h at RT. The organoids were then frozen in Tissue-Tek O.C.T. Compound (Sakura Finetek Europe, Alphen aan den Rijn, The Netherlands) and stored at -20°C until further processing. Cryosections were cut at 8 µm thickness using a Leica CM1900 cryostat (Leica Microsystems, Wetzlar, Germany) and transferred to glass slides. Immunofluorescence analysis was performed as previously described [41]. Sections were imaged on a Leica TCS SP8 confocal microscope (Leica Microsystems, Wetzlar, Germany) using the Leica Application suite X (v3.7.0.20979). Details about the antibodies used in this study are available in Table S4.

### Quantification and Statistical analysis

For quantification purposes, three representative images per organoid were acquired at 40X magnification. Quantification was performed using Fiji ImageJ (v2.17). Graphs were generated on GraphPad Prism (v10.2.3), where each data point corresponds to the mean per organoid ± standard error of the mean. The number of organoids tested is indicated in the figure legends. Experiments were performed on three independent rounds of differentiation. Normality was assessed using the Shapiro–Wilk test supported by visual inspection of data distribution via QQ plots. Since data violated normality assumptions, group differences were assessed using the Kruskal–Wallis test followed by Dunn’s multiple comparisons test. Significance was indicated as *p* < 0.05 (*), *p* < 0.01 (**), *p* < 0.001 (***), and ns (not significant).

### Single-Cell RNA Sequencing

Single-cell RNA sequencing was performed using the 10X Genomics Chromium 3′ gene expression platform. Organoids from a single differentiation batch were analyzed (ISO-CTRL n = 4; *ABCA4*-H10 *n* = 5; *ABCA4*-G3 *n* = 5). Individual organoids were dissociated into single-cell suspensions using the Papain Dissociation kit (Worthington Biochemical Corp., Lakewood, NJ, USA) following manufacturer’s instructions, and processed with Chromium v3 chemistry. Sequencing data were processed using Cell Ranger (v7.1.0) with the GRCh38 human reference genome. Downstream analyses were conducted in R (v4.4.0) using a Seurat-based pipeline. Differential gene expression was evaluated at the single-cell level using Wilcoxon rank-sum test, with multiple-testing correction performed using the Benjamini– Hochberg method. Genes with an adjusted p-value < 0.05 were considered significantly differentially expressed. Additional analytical details are provided in the Supplementary Methods.

## Results

### Generation of CRISPR/Cas9-engineered hiPSCs harboring a premature stop codon in exon-24 of ABCA4

Here, we applied the CRISPR/Cas9 gene editing tool to a previously characterized parental cell line, LUMC0004iCTRL10 [40], renamed as “ISO-CTRL”, to generate homozygous mutant hiPSC lines where a 10-bp deletion-insertion results in the introduction of an in-frame premature stop codon in *ABCA4* exon-24, c.3677-3686delinsTGA; p.(Val1192Ter) (Figure 1A). Two independent mutant hiPSC subclones, labeled as *ABCA4*-G3 and *ABCA4*-H10, were analyzed by Sanger sequencing to confirm the presence of the mutation (Figure 1B). Based on the structural insights into ABCA4 [42], this variant is predicted to generate a truncated ABCA4 protein within the regulatory domain 1 (R1; amino acids 1155-1279), located downstream of the nucleotide-binding domain 1 (NBD1; amino acids 915-1152 aa). G-banding indicated normal karyotypes (Figure S1A), and genome copy number variation analysis did not detect frequently occurring chromosomal abnormalities (Figure S1B), suggesting absence of genetic aberrations in the *ABCA4*-G3 and *ABCA4*-H10 lines. Lastly, the short tandem repeats analysis revealed that the mutant clones indeed share their derivation from the isogenic control line (Figure S1C). To rule out the occurrence of off-target mutagenesis, we used the IDT software “CRISPR-Cas9 guide RNA design checker” prediction tool (https://eu.idtdna.com/) to identify the top 5 candidate off-target sites and confirmed the absence of indels in the genomic regions of the *ABCA4* subclones by Sanger sequencing (Figure S2).

**Figure 1:**
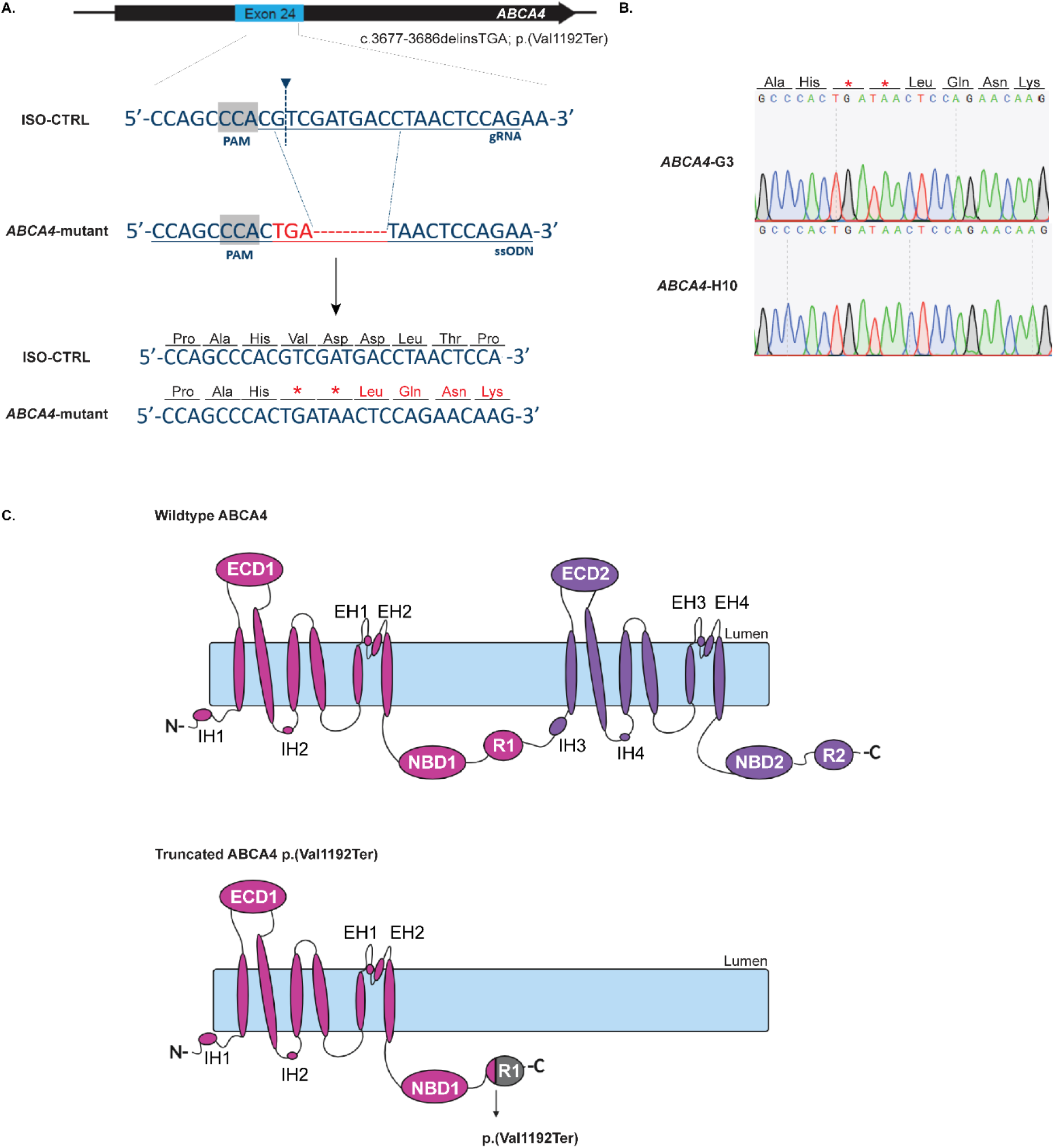
Generation of *ABCA4* hiPSCs and protein structure prediction. (A) Schematics indicating the gene disruption design, with a gRNA targeting exon-24 of *ABCA4*, and a single-stranded oligodeoxynucleotide (ssODN) donor inducing the deletion of 10 bp and the insertion of the premature stop codon TGA. (B) Sanger sequencing of the region of interest in exon-24 confirming the introduction of the mutation in homozygous form in *ABCA4* hiPSCs. (C) Depiction of the predicted protein domains in wildtype ABCA4 (top) [42] and truncated ABCA4 p.(Val1192Ter). Elongated ovals indicate transmembrane domains; ECD, extracellular domain; NBD, nucleotide-binding domain; R, regulatory domain; IH, intracellular helix; EH, extracellular helix. NBD1 domain: 915-1152 aa; R1 domain: 1155-1279 aa.

### ABCA4 hiPSC-derived retinal organoids show correct development of rod and cone photoreceptor cells, but loss of ABCA4 protein in OS-like structures

After validation of the hiPSC subclones, the *ABCA4*-G3 and *ABCA4*-H10 cell lines were differentiated into retinal organoids, along with their isogenic parental line, until differentiation day (DD) 240, following a protocol earlier established by our team [43]. The brightfield images showed a comparable development of the retinal lamination in both isogenic control and mutant organoids, with the formation of a characteristic brush border corresponding to the photoreceptor layer (Figure 2A). We did not observe morphological changes in the *ABCA4*-G3 and *ABCA4*-H10 retinal organoids that would be indicative of retinal degeneration.

**Figure 2:**
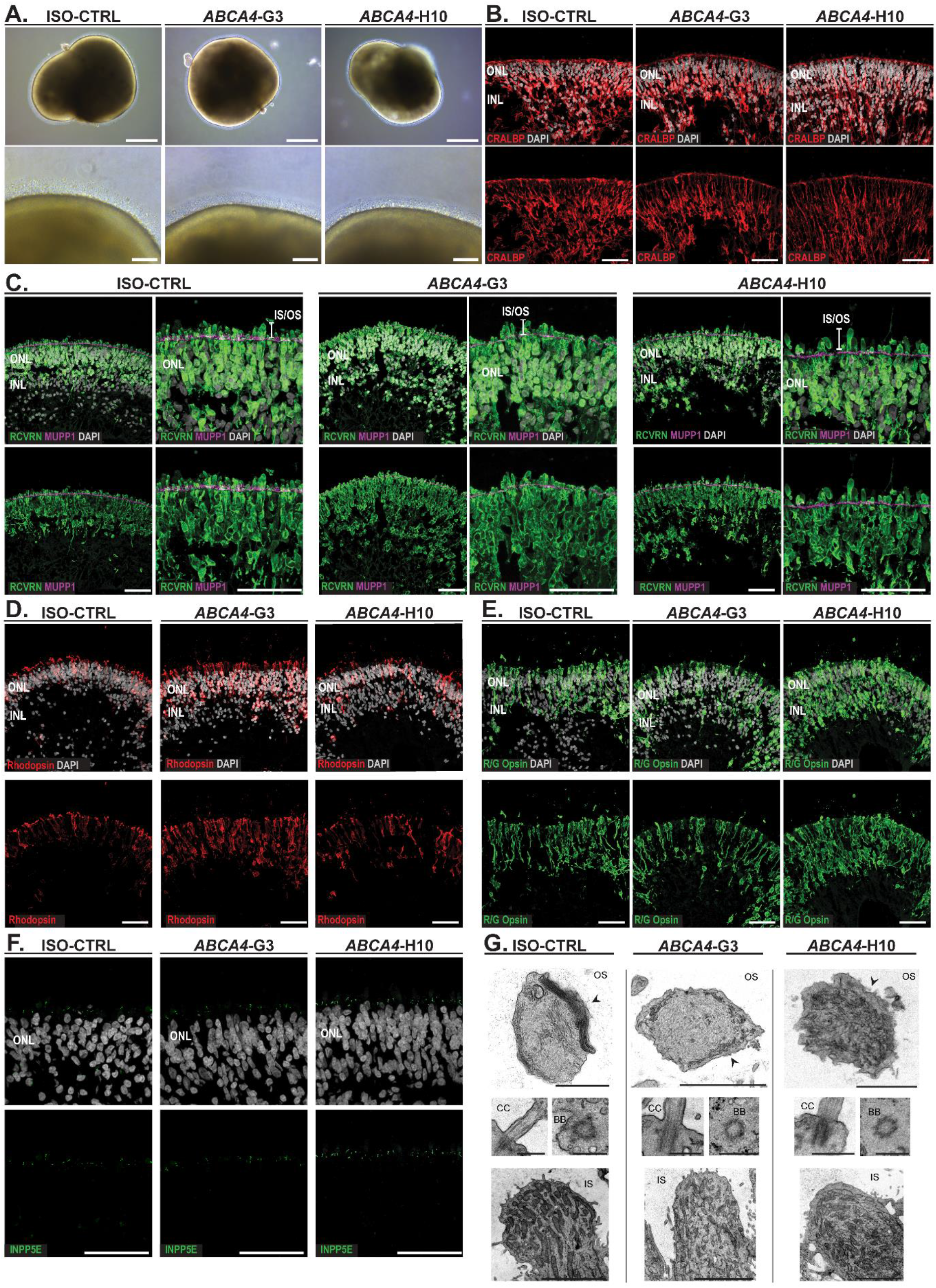
*ABCA4* hiPSC-derived retinal organoids show correct development of major retinal cell types at DD240. (A) Representative brightfield images of ISO-CTRL, *ABCA4*-G3 and *ABCA4*-H10 retinal organoids at DD240. Bottom images display the typical photoreceptor brush border at higher magnification. Scale bars: top, 500 µm; bottom, 100 µm. (B) Representative immunofluorescence Z-stack images of CRALBP (red) with and without DAPI (grey). Scale bars: 20 µm. (B-F) ONL = outer nuclear layer, INL = inner nuclear layer. (C) Representative immunofluorescence Z-stack images of recoverin (green) and MUPP1 (purple), with and without DAPI (grey). Photoreceptor inner segment = IS; photoreceptor outer segment (OS). Scale bars: left, 20 µm; right: 50 µm. (D) Representative immunofluorescence Z-stack images of rhodopsin (red), with and without DAPI (grey). Scale bars: 20 µm. (E) Representative immunofluorescence Z-stack images of red/green (R/G) opsin (green), with and without DAPI (grey). Scale bars: 20 µm. (F) Representative immunofluorescence Z-stack images of INPP5E (green), with and without DAPI. Scale bars: 50 µm. (E) Representative transmission electron microscopy (TEM) images of photoreceptors in ISO-CTRL and *ABCA4* retinal organoids at DD240. Upper panel: outer segment (OS)-like structures with arrows pointing to the disc membranes. Scale bars from left to right: 1 µm, 2 µm, 1 µm. Middle panel: connecting cilium (CC) and basal body (BB). Scale bars: 0.5 µm. Bottom panel: inner segment (IS)-like structures. Scale bars: 3 µm.

We previously demonstrated that organoids cultured using the same differentiation protocol as used in this study show correct development of all major retinal cell types, including rod and cone photoreceptor cells, Müller glial cells, ON and OFF bipolar cells and horizontal cells [39]. To confirm the appropriate differentiation of the *ABCA4* mutant hiPSC lines into retinal organoids, we performed immunofluorescence analysis using antibodies against key markers of selected cell types. CRALBP staining highlighted the presence of Müller glial cells, displaying the characteristic elongated, radial morphology, with filamentous processes extending from the inner retina toward their apical end feet (Figure 2B). Anti-recoverin marked the photoreceptor cells, with multiple PDZ domain protein 1 (MUPP1) antibody delineating the outer limiting membrane (OLM), allowing clear visualization of the inner (IS) and outer segments (OS) extending beyond it (Figure 2C). Ultimately, rods and cones were separately labelled by using antibodies against rhodopsin and R/G opsin, respectively, which are normally trafficked from the IS to the OS (Figure 2D, E). These two portions of the photoreceptor cells are known to be linked by the connecting cilium (CC), here detected in both control and *ABCA4* organoids via the inositol polyphosphate-5-phosphatase E (INPP5E) marker (Figure 2F). A deeper focus on the photoreceptor structure was provided by the transmission electron microscopy (TEM) analysis, performed according to an established protocol [45]. The images showed the presence of the photoreceptor inner segment-like regions containing mitochondria, the connecting cilium (CC) and related basal body (BB), and very immature outer segment (OS)-like structures, characterized by the presence of yet poorly organized photoreceptor discs (Figure 2G). Overall, qualitative comparison between isogenic control and *ABCA4* mutant organoids did not reveal overt differences, suggesting that *ABCA4* mutation does not affect retinal organoid development.

Due to the expected localization of the ABCA4 protein to the discs of rods and cones, we focused on further characterizing the outer segment of photoreceptor cells. We used two alternative disc markers, namely the rod outer segment membrane protein 1 (ROM1) and peripherin-2 (PHRP2), to investigate any potential changes to the ABCA4 location. Interestingly, while at DD240 ABCA4 was mostly co-localizing with ROM1 (Figure 3A), and PRPH2 (Figure 3B) in the isogenic control organoids, the mutant photoreceptors of *ABCA4*-G3 and *ABCA4*-H10 displayed loss of the ABCA4 protein at the outer segments. The anti-ABCA4 raised against the amino-terminal half of ABCA4 (amino acids 1-249) did not detect the truncated protein, suggesting that this shorter ABCA4 is degraded. A quantitative analysis performed on organoids from three independent batches of differentiation revealed no detectable changes in the intensity of ROM1 (Figure 3D) and PRPH2 (Figure 3E) in the photoreceptor cells of subclones *ABCA4*-G3 and *ABCA4*-H10, yet a significant decrease of the ABCA4 signal when compared to the ISO-CTRL (Figure 3C). These results validate the genetic ablation at protein level and further support that ABCA4 deficiency does not impair the expression of key markers of retinal organoids and, specifically, of photoreceptor outer segments.

**Figure 3:**
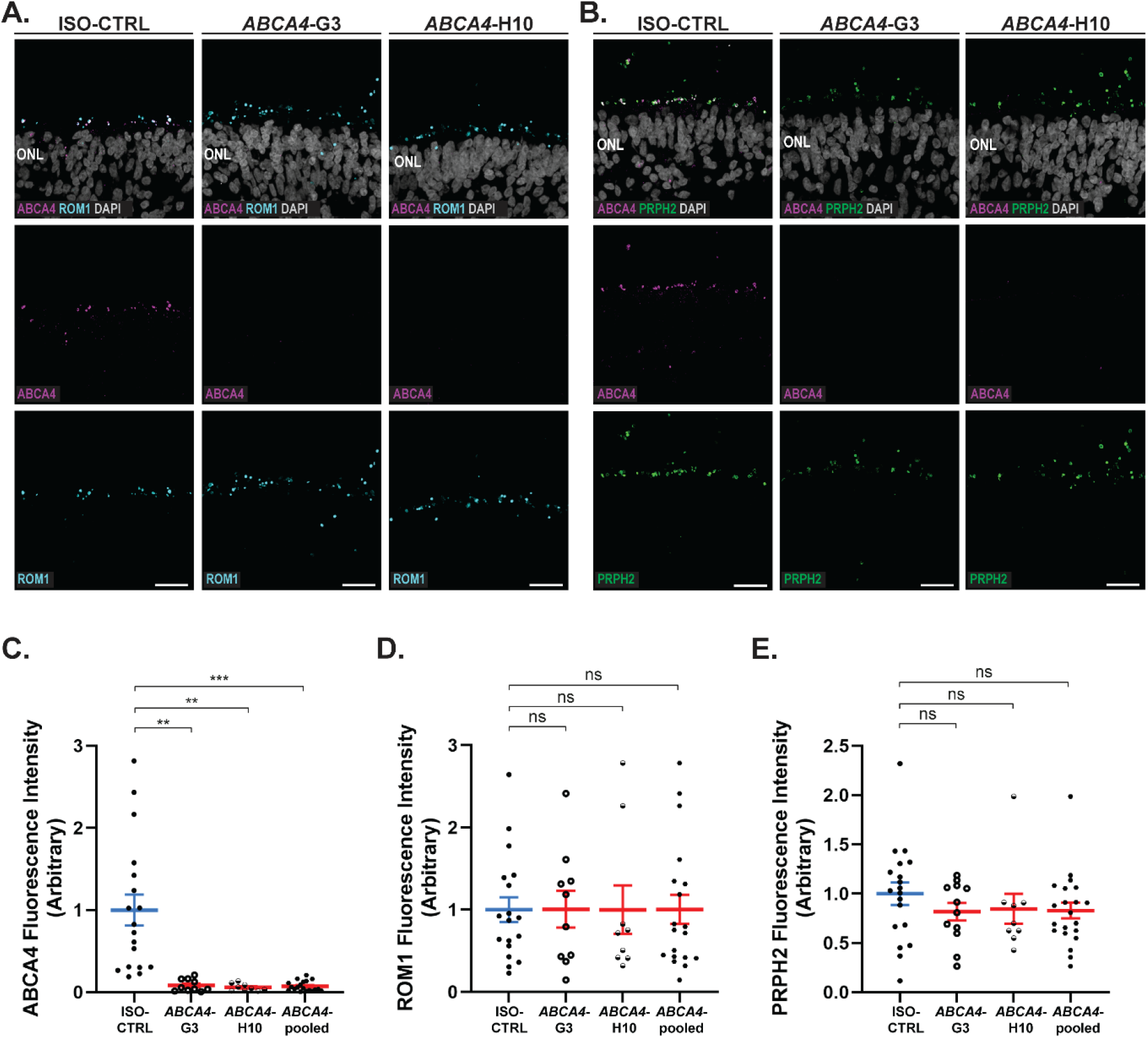
Loss of ABCA4 from the photoreceptor discs does not affect the localization of other markers in *ABCA4* retinal organoids at DD240. (A) Representative immunofluorescence images of ABCA4 (magenta) and ROM1 (cyan), with and without DAPI (grey), in ISO-CTRL, *ABCA4*-G3, and *ABCA4*-H10 retinal organoids at DD240. Scale bars: 20 µm. (B) Representative immunofluorescence images of ABCA4 (purple) and PRPH2 (green), with and without DAPI (grey), in ISO-CTRL, *ABCA4*-G3, and *ABCA4*-H10 retinal organoids at DD240. Scale bars: 20 µm. (C) Quantitative analysis of ABCA4 intensity in ISO-CTRL and *ABCA4* retinal organoids. Number of organoids used: ISO-CTRL *n* = 18; *ABCA4*-H10 *n* = 9; *ABCA4*-G3 *n* = 10, from three rounds of differentiation. Group differences were assessed using the Kruskal-Wallis test followed by Dunn’s multiple comparisons test. Significance was indicated as *p* < 0.01 (**), and *p* < 0.001 (***). Statistical analysis: *p* = 0.0097, *p* = 0.0010, *p* = 0.0003, from left to right. (D) Quantitative analysis of ROM1 intensity in ISO-CTRL and *ABCA4* retinal organoids. Number of organoids used: ISO-CTRL *n* = 18; *ABCA4*-H10 *n* = 9; *ABCA4*-G3 *n* = 10, from three rounds of differentiation. Group differences were assessed using the Kruskal-Wallis test followed by Dunn’s multiple comparisons test. Significance was indicated as ns (not significant). Statistical analysis: *p* > 0.9999 for all groups. (E) Quantitative analysis of PRPH2 intensity in ISO-CTRL and *ABCA4* retinal organoids. Number of organoids used: ISO-CTRL *n* = 19; *ABCA4*-H10 *n* = 9; *ABCA4*-G3 *n* = 12, from three rounds of differentiation. Group differences were assessed using the Kruskal-Wallis test followed by Dunn’s multiple comparisons test. Significance was indicated as ns (not significant). Statistical analysis: *p* = 0.9231, *p* = 0.3875, *p* = 0.3932, from left to right.

### Retention of ABCA4 transcripts in ABCA4 retinal organoids

To investigate potential transcriptomic alterations not associated with morphological defects, we performed single-cell RNA sequencing (scRNA-Seq) on control and *ABCA4* retinal organoids at DD225 (ISO-CTRL n = 4, *ABCA*4-G3 n = 5, *ABCA4*-H10 n =5). By performing 14 scRNA-Seq experiments, we identified 19 distinct clusters displayed on a UMAP plot (Figure 4A), including rod and cone photoreceptors, Müller glial cells, ON and OFF bipolar cells, horizontal cells, and amacrine cells. Cluster identities were assigned based on established marker gene expression (Figure 4B) [46, 47]. Unexpectedly, several additional cell types were identified compared to the clusters found in a previous analysis conducted on organoids differentiated using the same protocol [39]. Such cells were accounted for as astrocytes, dividing cells, fibroblasts, and multiple transient cell populations. These transient clusters comprised bipolar–photoreceptor cells (T1), cone–bipolar–horizontal cells (T2), MGC–cone cells (T3), and bipolar–amacrine–horizontal cells (T4). Multiple MGC clusters were detected and annotated as MGCs-I–IV. Cell-type proportions were comparable across ISO-CTRL and *ABCA4* organoids, indicating no major differences in cellular composition, further suggesting that loss of ABCA4 does not significantly impact retinal organoid development (Figure 4C).

**Figure 4:**
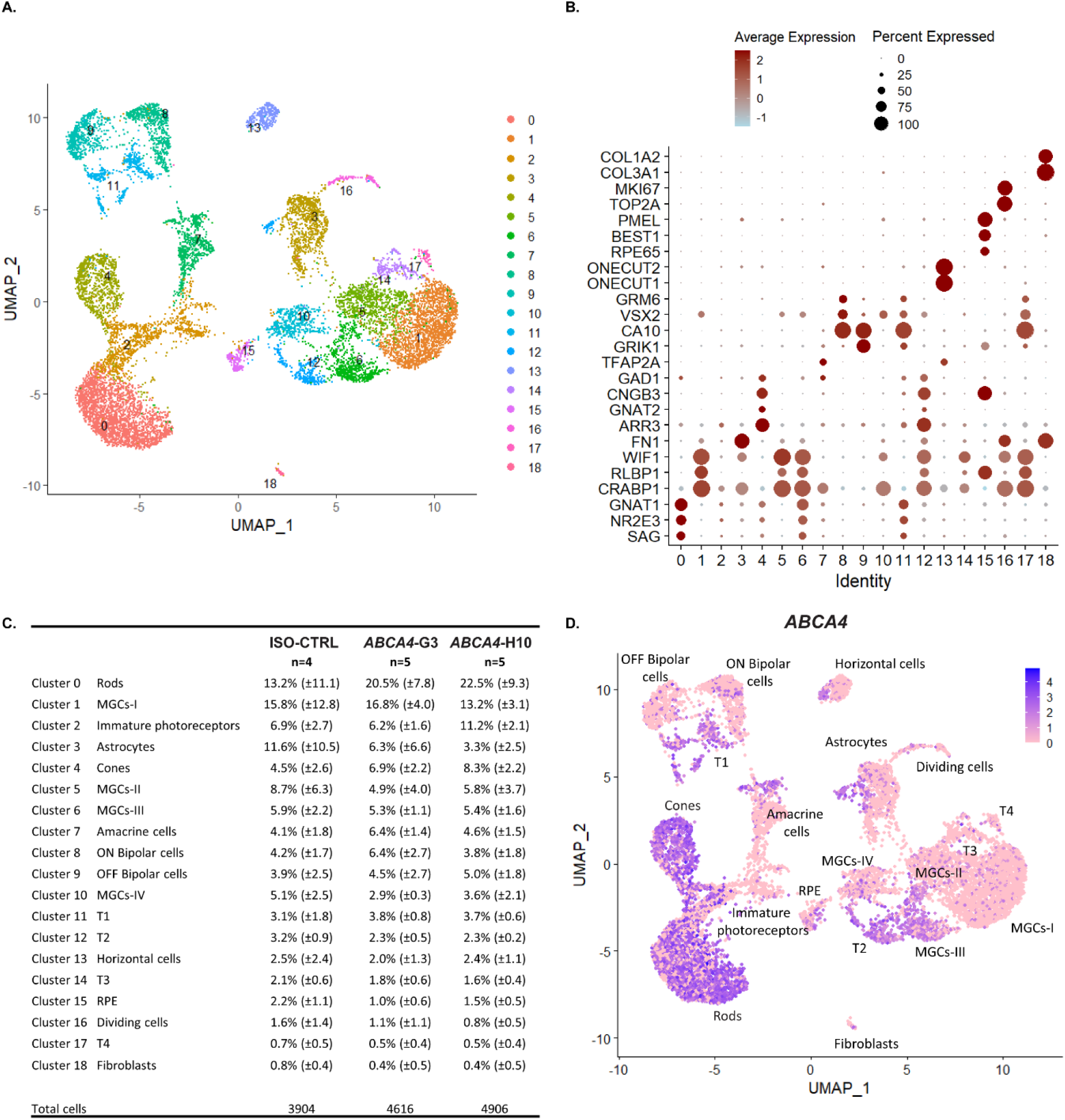
Cell-type identification and *ABCA4* transcript enrichment at DD225 in hiPSC-derived *ABCA4* and control retinal organoids by single-cell RNA sequencing. (A) UMAP displaying 19 distinct clusters from 14 scRNA-Seq experiments. (B) Dot plot showing transcript enrichment of cell-specific markers to assign cell identity. (C) Table indicating the percentage of each cell type across conditions including standard deviation. (D) UMAP plot showing *ABCA4* transcript enrichment. Number of organoids used: ISO-CTRL *n* = 4; *ABCA4*-H10 *n* = 5; *ABCA4*-G3 *n* = 5 from 14 scRNA-Seq of the same differentiation round.

Once that all cell identities were attributed, we proceeded to analyze the expression of *ABCA4* in the different cell types. *ABCA4* transcripts were found to be predominantly enriched in photoreceptor cells and transient populations, with lower expression levels detected in a subset of MGCs, amacrine cells, horizontal cells, astrocytes, and RPE (Figure 4D).

Although a premature stop codon was introduced in exon-24 of *ABCA4* and expected to activate the nonsense-mediated mRNA decay, no statistically significant reduction in *ABCA4* transcript levels was detected in most cell types of *ABCA4* organoids (Figure S3A, S3D, S3G-I). The sole exception was the rod photoreceptors from the *ABCA4*-H10 subclone, where *ABCA4* was identified as a differentially expressed gene (DEG) (Figure S3A). Consistent with the immunofluorescence data showing preserved localization of ROM1 and PRPH2 proteins, transcript levels of these disc markers did not change in both rod and cone photoreceptors of *ABCA4* organoids (Figure S3B, S3C, S3E, S3F).

Differential expression analysis revealed minimal transcriptional changes in the photoreceptor populations. Rods displayed a limited number of DEGs, only 5 in *ABCA4*-G3 (Figure S4A) and 37 genes in *ABCA4*-H10 (Figure S4B). Among these, *B2M* and the long non-coding RNA *AL451062*.*1* were the only genes consistently dysregulated across both *ABCA4* subclones. On the other hand, cone photoreceptors showed no DEGs in *ABCA4*-G3 (Figure S4C) and only three DEGs in *ABCA4*-H10 (*NRL, CC2D2A*, and *B2M*) (Figure S4D). These findings indicate that, despite the loss of ABCA4 protein from the photoreceptor discs, rods and cones in *ABCA4* hiPSC-derived retinal organoids largely retain a stable transcriptomic profile at DD225.

### scRNA-Seq analysis reveals major transcriptional response in ABCA4 Müller glial cells and astrocytes

In contrast to photoreceptors, a substantial transcriptional response was observed in Müller glial cells and astrocytes. In the MGCs-I cluster, *ABCA4*-G3 (Figure S5A) and *ABCA4*-H10 (Figure S5B) displayed 70 and 317 DEGs, respectively, with 29 genes commonly dysregulated across subclones. These included 13 upregulated genes, such as *SAT1, PHLDA1, FABP5*, and *SQSTM1* (Figure 5Ai), and 16 downregulated genes, including *TF, PCDHGA3, BNIP3*, and *HEY1* (Figure 5Aii). Similar expression patterns were observed across additional MGC clusters, with consistent downregulation of *TF* in MGCs-I, MGCs-II, and MGCs-III, and downregulation of *CRYM* in MGCs-I and MGCs-II. We also detected upregulation of glial markers such as *CD44* in MGCs-I and MGCs-III, and upregulation of *FTL* in MGCs-I, MGCs-II, MGCs-III, MGCs-IV and T2. *IFITM3* was found upregulated in MGCs-II and MGCs-IV. In cluster MGCs-IV other interferon-induced genes as well were found upregulated, namely *IFITM1* and *IFITM2*.

**Figure 5:**
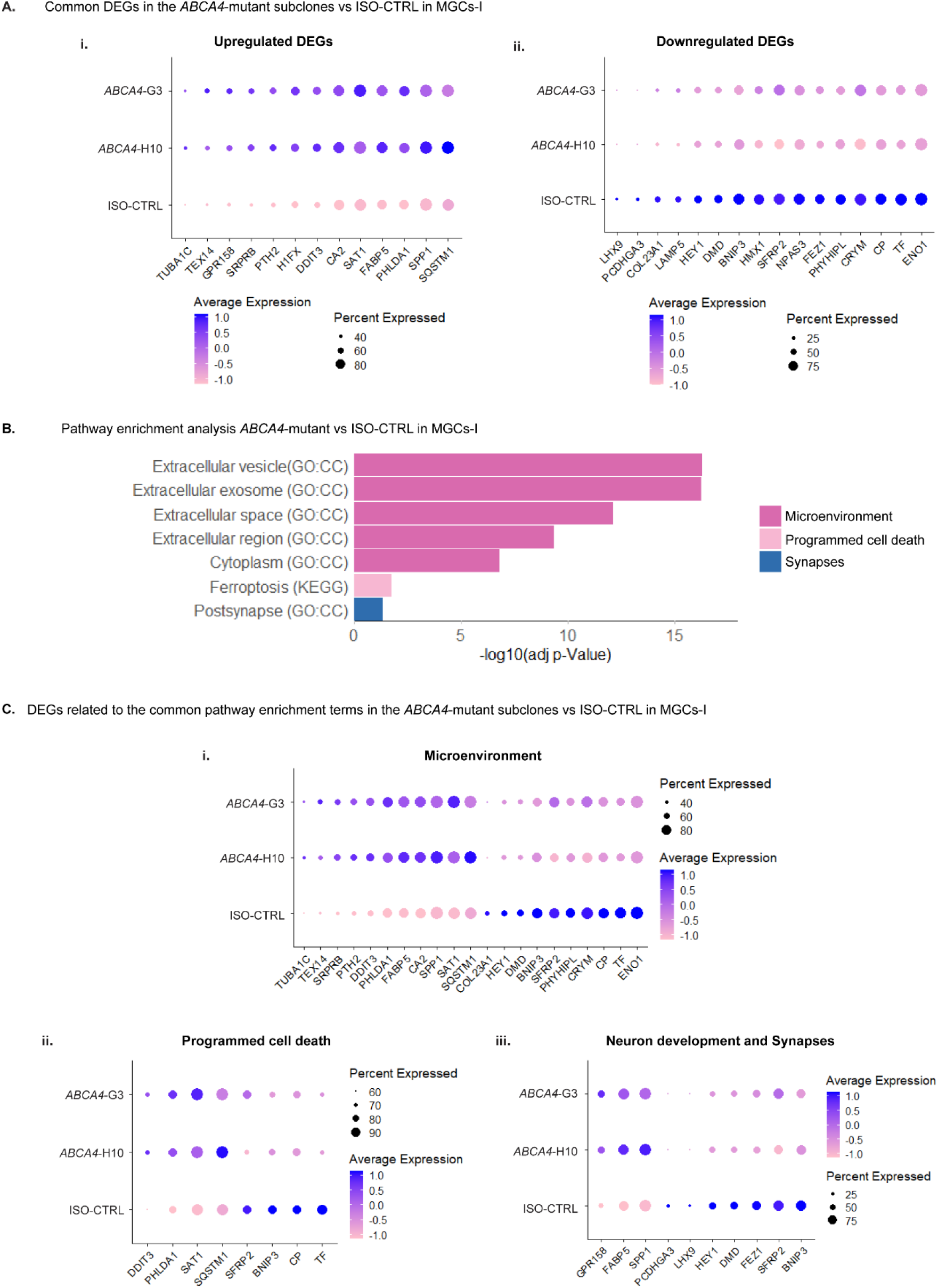
scRNA-Seq analysis shows a transcriptional response of Müller glial cells upon *ABCA4* mutation. (A) Pathway enrichment analysis of differentially expressed genes (DEGs) in MGCs-I, with similar terms grouped into three categories. (B) Dot plots of the statistically significant 13 upregulated DEGs (i) and 16 downregulated DEGs (ii) found in the MGCs of both *ABCA4* subclones. (C) Common DEGs involved in the shared pathway enrichment terms related to microenvironment (i), programmed cell death (ii), and neuron development and synapses (iii). Number of organoids used: ISO-CTRL *n* = 4; *ABCA4*-H10 *n* = 5; *ABCA4*-G3 *n* = 5 from 14 scRNA-Seq of the same differentiation round. Differential gene expression was evaluated at the single-cell level using Wilcoxon rank-sum test, with multiple-testing correction performed using the Benjamini–Hochberg method. Genes with an adjusted p-value < 0.05 were considered significantly differentially expressed.

*PCDHGA3*, encoding a small calcium-dependent neural cadherin-like cell adhesion protein, was markedly downregulated in all the *ABCA4* mutant four MGC clusters, along with astrocytes, amacrine cells, and the transient cluster T2, despite at low-level of expression. In some of these clusters, a complete loss of *PCDHGA3* expression was observed, suggesting impaired intercellular communication within the retinal network.

Pathway enrichment analysis of MGCs-I revealed enrichment in terms related to “microenvironment”, such as extracellular vesicle and cytoplasm, “programmed cell death”, and particularly ferroptosis, and “synapses”, with an accent on the postsynaptic specialization (Figure 5B). We then investigated which of the common DEGs found in the MGCs-I cluster of *ABCA4*-G3 and *ABCA4*-H10 were associated to such categories and displayed them in dot plots (Figure 5C). For the microenvironment group, multiple DEGs were retrieved, including *FABP5, COL23A1, SFRP2*, and *CRYM* (Figure 5Ci), whereas enrichment of programmed cell death was driven by *PHLDA1, BNIP3* (Figure 5Cii). The post-synapse, considered one of the later stages of the broader process of neuronal development, was associated with DEGs such as *SPP1, LHX9*, and *FEZ1* (Figure 5Ciii). Notably, *PCDHGA3* was also linked to this category and was consistently downregulated.

Astrocytes exhibited the strongest transcriptional alteration, with 113 DEGs in *ABCA4*-G3 (Figure S5C) and 447 DEGs in *ABCA4*-H10 (Figure S5D), of which 66 genes were shared between subclones (Figure 6A). These common DEGs included *PHLDA1, ADGRV1, BNIP3L*, and *NR2E1*, functionally overlapping with those identified in the MGCs. A similar pathway enrichment analysis performed on the astrocytes cluster revealed enrichment in terms related to “neuron development”, “cell-cell interaction”, “microenvironment”, and “programmed cell death” (Figure 6B). Among the DEGs contributing to neuronal development pathways, *NEGR1, NR2E1, NRG1* and *ADGRV1* were downregulated (Figure 7A). *PCDHGA3* was again found to be downregulated, implicating impaired cell-cell interactions (Figure 7B). Enrichment of microenvironment-related terms involved genes such as *COL18A1* and *APOC1* (Figure 7C), whereas *PHLDA1* and *BNIP3L* contributed to the dysregulation of programmed cell death (Figure 7D).

**Figure 6:**
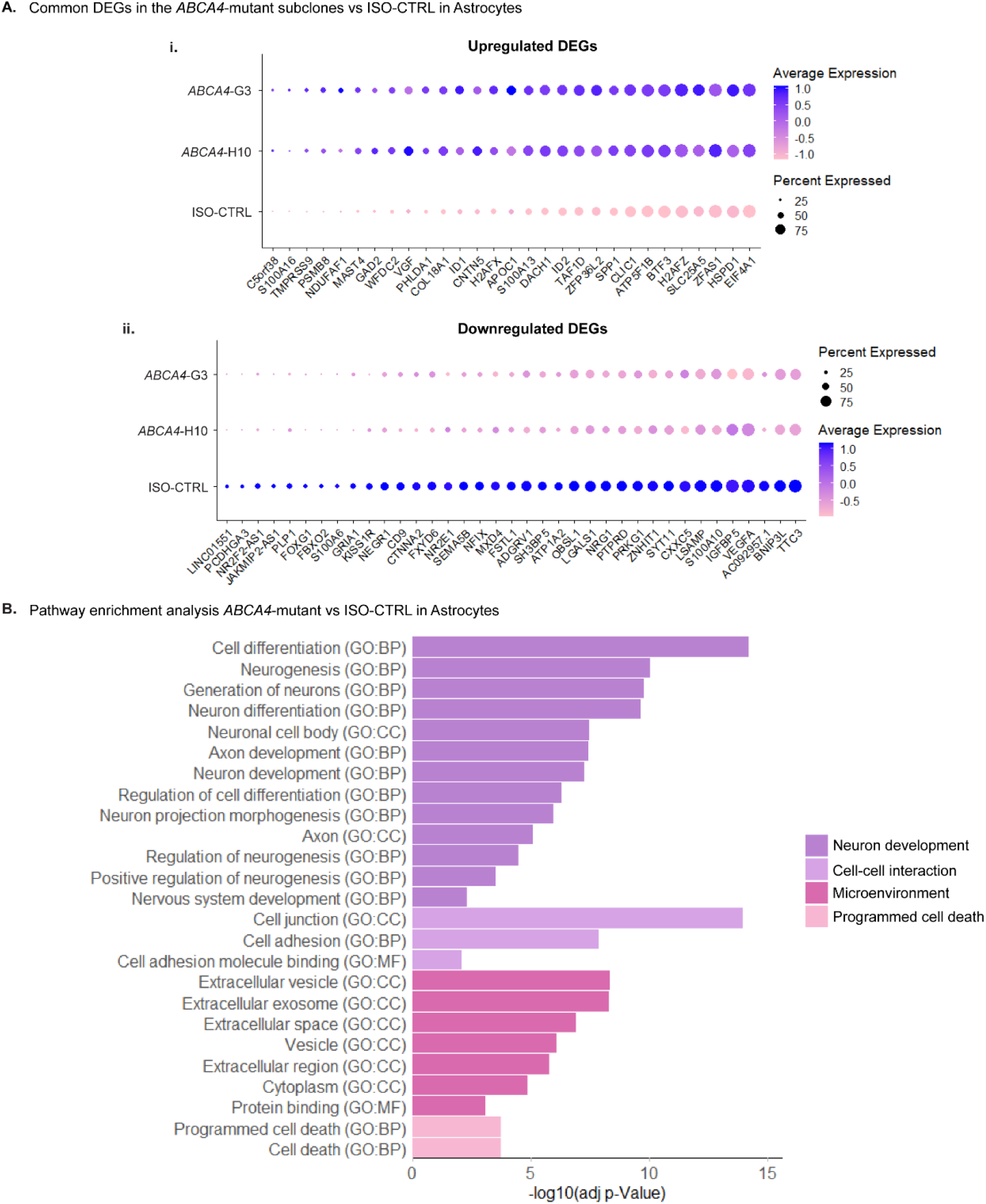
Astrocytes transcriptional profile is strongly modified upon *ABCA4* gene disruption. (A) Pathway enrichment analysis of differentially expressed genes (DEGs) in astrocytes, with similar terms grouped into four categories. (B) Dot plots of the statistically significant 29 upregulated DEGs (i) and 37 downregulated DEGs (ii) found in the astrocytes of both *ABCA4* subclones. Number of organoids used: ISO-CTRL *n* = 4; *ABCA4*-H10 *n* = 5; *ABCA4*-G3 *n* = 5 from 14 scRNA-Seq of the same differentiation round. Differential gene expression was evaluated at the single-cell level using Wilcoxon rank-sum test, with multiple-testing correction performed using the Benjamini–Hochberg method. Genes with an adjusted p-value < 0.05 were considered significantly differentially expressed.

**Figure 7:**
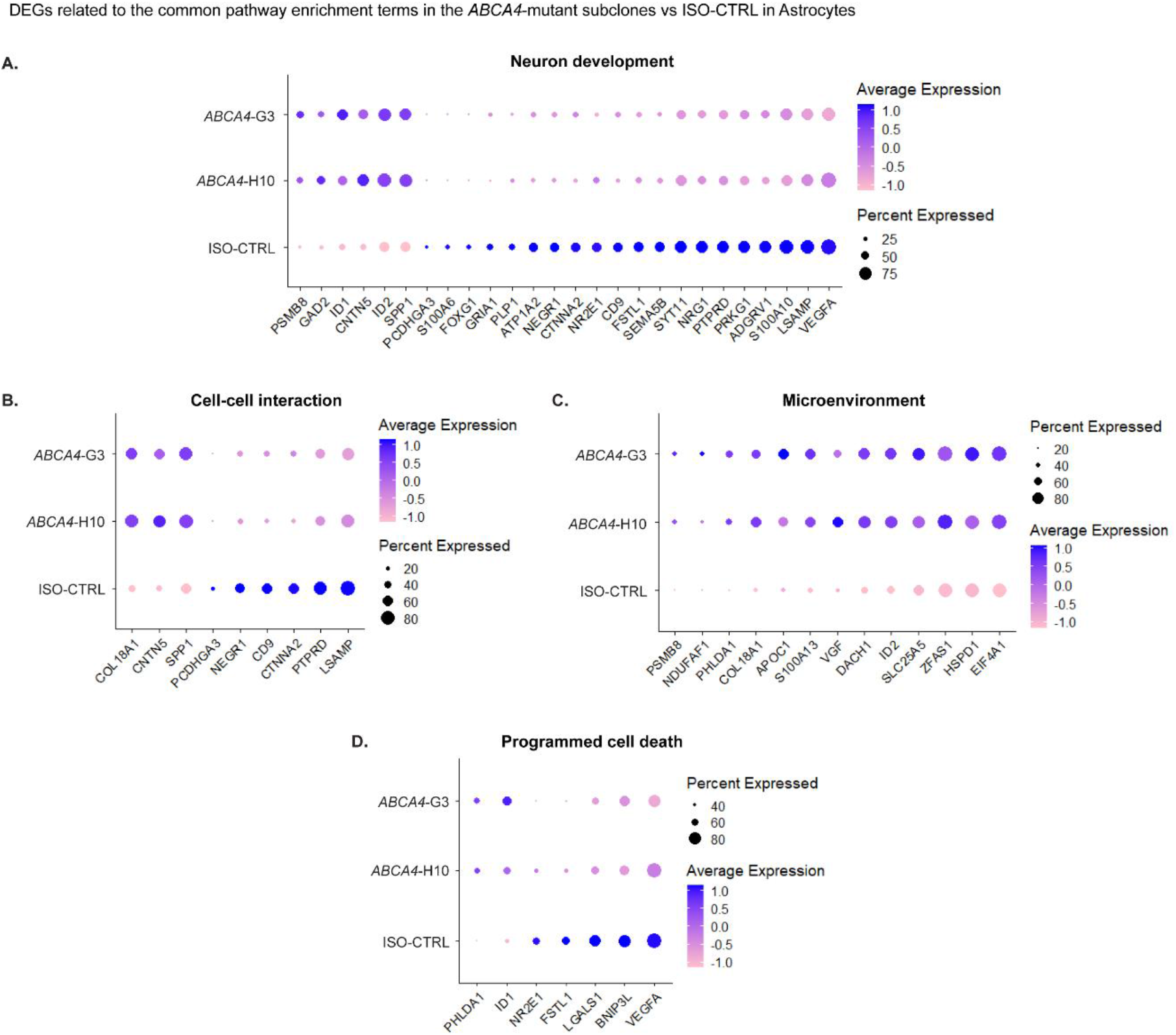
Differential expression analysis of *ABCA4* mutant astrocytes compared to isogenic controls. (A-D) Common DEGs involved in the shared pathway enrichment terms related to neuron development (A), cell-cell interaction (B), microenvironment (C), and programmed cell death (D). Number of organoids used: ISO-CTRL *n* = 4; *ABCA4*-H10 *n* = 5; *ABCA4*-G3 *n* = 5 from 14 scRNA-Seq of the same differentiation round. Differential gene expression was evaluated at the single-cell level using Wilcoxon rank-sum test, with multiple-testing correction performed using the Benjamini–Hochberg method. Genes with an adjusted p-value < 0.05 were considered significantly differentially expressed.

While our primary focus was on shared pathway alterations reflecting consistent and reproducible effects of ABCA4 loss, clonal variability must be taken into account, given the well-known heterogeneity of organoids. In fact, individual clones may manifest the phenotype differently, with some exhibiting a stronger response to the mutation, and others being more tolerant. When extending the pathway enrichment analysis to include terms uniquely enriched in one *ABCA4* subclone, but not in the other, additional subclone-specific biological pathways emerged. In the MGCs-I cluster of *ABCA4*-H10 organoids, we identified terms related to “cell-cell interaction”, “neuron development”, “iron binding and transport” and “cell redox homeostasis” (Figure S6A). Moreover, the “programmed cell death” category expanded to include pathways related not only to ferroptosis but also to apoptosis, another form of regulated cell death. On the other hand, *ABCA4*-G3 showed enrichment in pathways related to the “respiratory chain” and unique molecular function terms associated with “retinoid binding” (Figure S6B). Genes contributing to these subclone-specific categories were retrieved and plotted, highlighting DEGs unique to either the *ABCA4*-G3 or *ABCA4*-H10 group (n = 5 each) (Figure S7). The enrichment of “cell redox homeostasis” and “respiratory chain” pathways in *ABCA4*-H10 and *ABCA4*-G3, respectively, suggests the presence of ongoing metabolic stress in both subclones.

Additional subclone-specific pathway enrichments were also observed in the astrocytes. Terms related to “synapses” and postsynaptic specialization were exclusively enriched in *ABCA4*-H10 organoids (Figure S8A), whereas *ABCA4*-G3 astrocytes displayed enrichment in “glycolysis” and, indirectly, in pathways linked to the “respiratory chain complex” (Figure S8B). Corresponding supplemental dot plots are provided in Figure S7.

## Discussion

The aim of this study was to investigate the effects of *ABCA4* gene disruption on transcript and protein expression in CRISPR/Cas9-engineered hiPSC lines differentiated into retinal organoids. Although the introduction of a premature stop codon in exon-24 was predicted to generate a truncated ABCA4 protein within the regulatory domain 1 (R1), ABCA4 was undetectable in the photoreceptor outer segment discs when using an antibody targeting the N-terminal half of the protein. The ABCA4 loss was accompanied by altered transcriptional signatures in Müller glial cells and astrocytes. Specifically, we observed a disruption of pathways involved in the neuronal development, glial cell microenvironment, and programmed cell death.

Immunofluorescence results indicated that, despite the homozygous gene disruption of *ABCA4*, mutant retinal organoids exhibited comparable expression of markers for the major retinal cell types to those observed in the isogenic controls. Overall, we did not detect morphological changes in the *ABCA4*-G3 and *ABCA4*-H10 retinal organoids, as expected since the major cause for retinal degeneration is the toxic effects of bisretinoid A2E accumulated into retinal pigment epithelium [4, 7, 44]. Ultrastructural analysis by TEM demonstrated preserved photoreceptor architecture in *ABCA4* organoids, including identifiable connecting cilia, basal bodies, and inner segment-like and outer segment (OS)-like structures. The mutation resulted in the loss of ABCA4 protein from the photoreceptor outer segment discs, without affecting the localization of other disc-associated markers, such as ROM1 and PRPH2, which co-localize with ABCA4 in the control organoids.

Unexpectedly, our scRNA sequencing analysis revealed that *ABCA4* transcript levels were not significantly reduced in mutant photoreceptors, with the exception of rods from the *ABCA4*-H10 subclone in which *ABCA4* was identified as a differentially expressed gene. The retention of *ABCA4* transcripts suggests that the nonsense-mediated decay mechanism is not efficient, leading to the translation of an unstable and rapidly degraded truncated protein. Overall, mutant rods and cones did not exhibit major transcriptional alterations, despite the protein being physiologically localized to the photoreceptor cells. In fact, a limited number of common DEGs was detected in these clusters: none in cones and only two shared dysregulated genes in *ABCA4* rods.

The use of retinal organoids to model Stargardt disease has been previously explored [27, 48-50]. In contrast to our results, Rui Chen and colleagues earlier described marked transcriptomic alterations in *ABCA4* patient-derived photoreceptors at DD300, identifying 2074 and 694 downregulated genes in cones and rods, respectively [48]. Notably, a considerable fraction of these DEGs was not restricted to photoreceptors, but was also detected in other retinal cell types. Importantly, their scRNA-Seq analysis compared patient-derived organoids to non-isogenic healthy controls, which may partly account for the discrepancies observed between their results and ours. Nevertheless, their gene ontology (GO) analysis revealed enrichment of apoptotic pathways and reduced neuronal functions, including processes related to development and microenvironment of glial cells, and intercellular communication in the organoids differentiated from patient hiPSCs carrying mutations in the *ABCA4* gene compared to unrelated controls. These observations are consistent with our findings in non-photoreceptor retinal populations, which displayed a pronounced transcriptional response to *ABCA4* deficiency.

In fact, in our study Müller glial cells exhibited 29 common DEGs, enriched in pathways related to disruption of glial microenvironment, regulation of programmed cell death, and multiple stages of neuronal development. Astrocytes showed an even stronger response, with 66 shared DEGs across subclones, involved in similar functional categories and additionally highlighting pathways associated with cell–cell interactions.

Among the dysregulated genes, *ADGRV1*, encoding adhesion G protein-coupled receptor V1, was downregulated in both Müller glial cells and astrocytes. While *ADGRV1* is known to participate in protein trafficking between the inner and outer segments of photoreceptors as part of the USH2 complex [51], it also plays a role in glutamate homeostasis in astrocytes, thereby supporting neuronal development [52]. Its downregulation in glial populations suggests that neuron–glia signaling and neuronal maturation may be indirectly affected by the ABCA4 protein loss, although the mechanistic link between *ABCA4* deficiency and *ADGRV1* regulation remains to be elucidated.

Another notable finding was the downregulation of *PCDHGA3*, a gene encoding a protocadherin involved in cell adhesion, whose ubiquitination modulates cell-cell interaction [53]. Indeed, this gene was retrieved from gene ontology terms indicative of intercellular communication and neuronal development. *PCDHGA3* expression was significantly reduced across multiple cell populations in *ABCA4* retinal organoids, including several Müller glial cell clusters, astrocytes, amacrine cells, and the transient T2 cluster. In some of these populations, *PCDHGA3* expression was completely ablated, suggesting a potential impairment in the establishment or maintenance of specific retinal cellular connections.

Programmed cell death also emerged as a shared dysregulated pathway in Müller glial cells and astrocytes. In addition to signatures related to iron-dependent ferroptosis, several GO terms associated with apoptotic pathways and their regulation were enriched. This may reflect alterations in the highly regulated cell death processes that physiologically occur during the retinal development. The upregulation of *PHLDA1*, identified as a DEG in both Müller glial cells and astrocytes, suggests a role in modulating these pathways. Conversely, *BNIP3* was downregulated in Müller glial cells, while *BNIP3L* was downregulated in astrocytes, further supporting cell-type-specific modulation of the apoptotic signaling.

Taken together, these results indicate that the *ABCA4* gene disruption primarily elicits a transcriptional response in Müller glial cells and astrocytes, rather than in photoreceptor cells. The loss of ABCA4 protein from the photoreceptor discs was not accompanied by major transcriptional alterations. Instead, the substantial transcriptional changes observed in Müller glial cells and astrocytes might represent an early response to *ABCA4* deficiency and/or subtle alterations in photoreceptor homeostasis, potentially reflecting a nonspecific reactive glial state (gliosis). This reactive response is characterized by altered expression of genes involved in neurodevelopmental processes and microenvironment of glial cells, highlighting the sensitivity of these cells to perturbations in the more resistant photoreceptors.

These observations align with our recent findings in *USH2A*-13KO hiPSC-derived retinal organoids, in which loss of usherin isoform B from the connecting cilium resulted in only mild transcriptional alterations in rod photoreceptors, but a markedly stronger disruption of homeostasis in Müller glial cells [39]. The involvement of MGCs in inherited retinal diseases is documented, highlighting their role as early responders to photoreceptor stress and their capacity to exhibit gene dysregulation prior to photoreceptor dysfunction [54]. Consistent with this, scRNA-Seq analysis of *Abca4* mice models has shown that mutant MGCs display early transcriptional responses to light-induced retinal degeneration, including dysregulation of synaptic connections and extracellular matrix remodeling [14].

*In vivo*, STGD1 pathology is associated to the transfer of photoreceptor-derived bisretinoids to the RPE through outer segment phagocytosis, leading to lysosomal dysfunction and oxidative stress in the RPE, and subsequent photoreceptor degeneration [44, 55]. However, in our organoid system, photoreceptors do not interface with a mature functional RPE monolayer. As a result, outer segment turnover and disc phagocytosis by RPE are absent, likely preventing the cascade that normally leads to the photoreceptor degeneration. This might explain why rods and cones are able to maintain a stable transcriptional profile despite the loss of ABCA4 protein.

The pronounced transcriptional alterations exhibited by MGCs and astrocytes would therefore represent an early, RPE-independent response that precedes photoreceptor death. Future studies should focus on the incorporation of a mature and functional retinal pigment epithelium into the organoid model. Such co-culture system would enable the physiological outer segment phagocytosis and retinoid exchange, allowing us to investigate whether the *ABCA4* deficiency leads to a secondary photoreceptor degeneration.

Furthermore, the analyses in this study were performed using two independent *ABCA4* subclones derived from the same parental line. While the use of isogenic controls reduces background genetic variability, basing conclusions on only two edited lines represents a limitation, as clone-specific effects cannot be entirely excluded. Follow-up experiments with additional independent mutant clones are essential to confirm the reproducibility of our findings, and specifically the transcriptomic data. In fact, while immunofluorescence results were validated across three differentiation rounds, scRNA-Seq analysis was performed on organoids from a single batch, leaving the possibility of batch-related effects.

Nevertheless, retinal organoids offer a relevant system for evaluating novel gene therapy approaches. Dual AAV strategies have emerged as a promising approach to overcome the packaging limitations associated with delivery of large genes, such as *ABCA4*. Intein-mediated protein trans-splicing has enabled efficient reconstitution of full-length ABCA4 protein in retinal cells [27], and has demonstrated both safety and therapeutic efficacy in preclinical animal models [26]. More recent advances, including mRNA trans-splicing [56] and AAV with translocation linkage [57], have further expanded the versatility and precision of large cargo delivery. In parallel, AAV-mediated homology-independent targeted integration (HITI) has been explored as an alternative strategy for retinal gene therapy, successfully ameliorating the phenotype of mouse models of retinitis pigmentosa [58]. Collectively, these studies highlight the rapid progress and growing therapeutic potential of gene therapies for inherited retinal disorders, including Stargardt disease. The *ABCA4* mutant retinal organoids here presented and characterized provide a relevant human model system to test the efficacy of *ABCA4* homology-independent targeted integration (HITI) gene therapy vectors.

## Conclusion

In conclusion, our data demonstrate that the introduction of a premature stop codon in exon-24 of the *ABCA4* gene results in the loss of ABCA4 protein from the photoreceptor outer segment discs, without inducing detectable transcriptional alterations in rods and cones. Unexpectedly, other retinal cell populations, particularly Müller glial cells and astrocytes, exhibited significant transcriptomic changes, affecting neuronal development, intercellular communication, and programmed cell death pathways. Further functional investigation is needed to assess the consequences of the perturbation of such pathways and to clarify whether MGCs and astrocytes contribute to the onset or progression of Stargardt disease.

## Acknowledgments

The authors thank their team members for valuable discussions and advice.

## Data Availability

The data underlying this article are available in the article and in its online supplementary material, except the scRNA-Seq data that are accessible at the GEO repository (GSE326273).”

