## Supplementary materials for "Loss of ABCA4 from photoreceptor discs triggers changes in glial cell homeostasis"

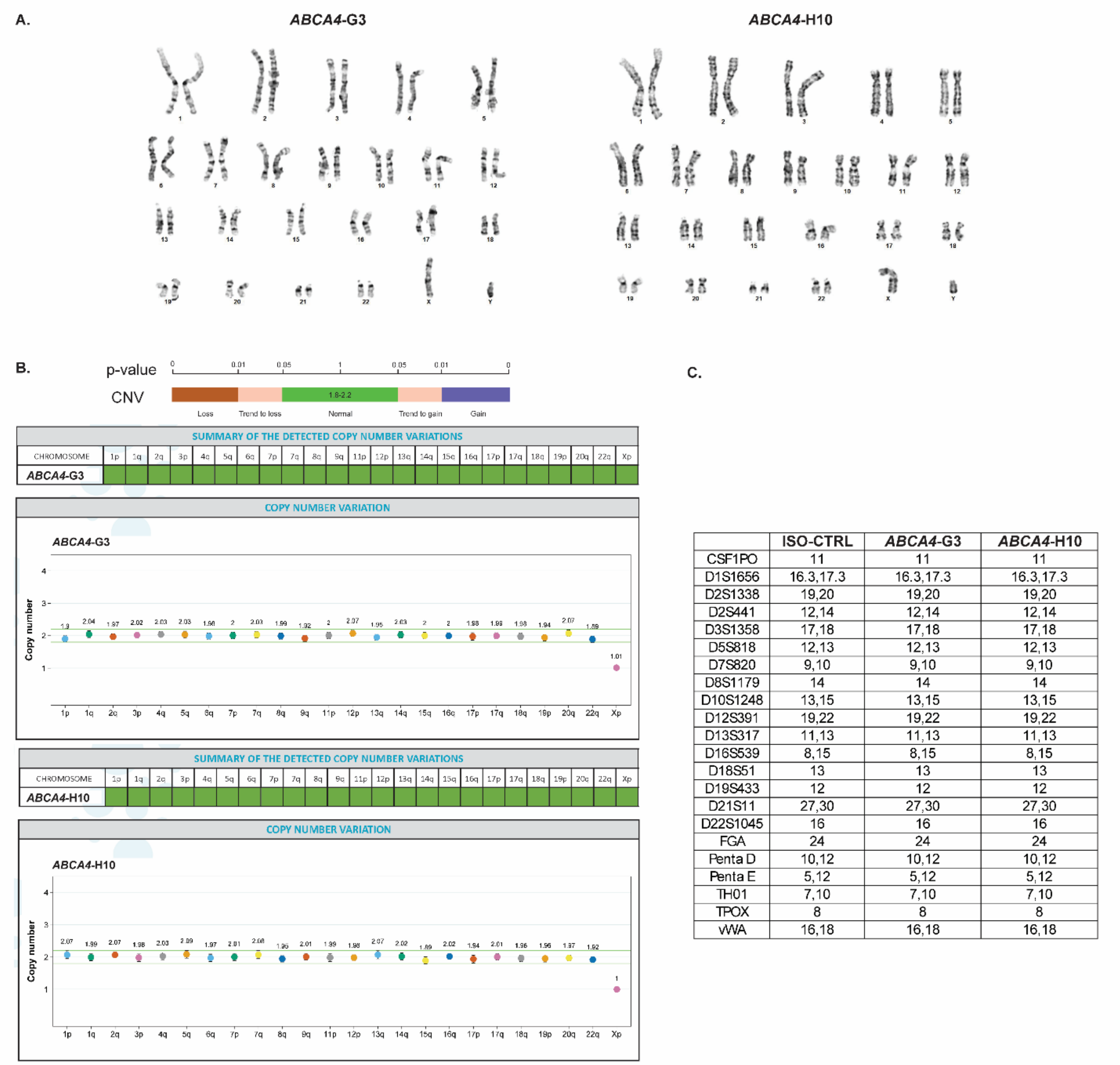


| | ISO-CTRL | *ABCA4*-G3 | *ABCA4*-H10 |
|---|---|---|---|
| CSF1PO | 11 | 11 | 11 |
| D1S1656 | 16.3,17.3 | 16.3,17.3 | 16.3,17.3 |
| D2S1338 | 19,20 | 19,20 | 19,20 |
| D2S441 | 12,14 | 12,14 | 12,14 |
| D3S1358 | 17,18 | 17,18 | 17,18 |
| D5S818 | 12,13 | 12,13 | 12,13 |
| D7S820 | 9,10 | 9,10 | 9,10 |
| D8S1179 | 14 | 14 | 14 |
| D10S1248 | 13,15 | 13,15 | 13,15 |
| D12S391 | 19,22 | 19,22 | 19,22 |
| D13S317 | 11,13 | 11,13 | 11,13 |
| D16S539 | 8,15 | 8,15 | 8,15 |
| D18S51 | 13 | 13 | 13 |
| D19S433 | 12 | 12 | 12 |
| D21S11 | 27,30 | 27,30 | 27,30 |
| D22S1045 | 16 | 16 | 16 |
| FGA | 24 | 24 | 24 |
| Penta D | 10,12 | 10,12 | 10,12 |
| Penta E | 5,12 | 5,12 | 5,12 |
| TH01 | 7,10 | 7,10 | 7,10 |
| TPOX | 8 | 8 | 8 |
| vWA | 16,18 | 16,18 | 16,18 |

**Figure S1: Validation of *ABCA4*-mutant hiPSCs.** (A) Karyotyping analysis showing normal metaphases. (B) Copy number variations (CNV) analysis indicating no abnormalities in the mutant subclones. (C) Short tandem repeats showing derivation of the *ABCA4* lines from the ISO-CTRL.

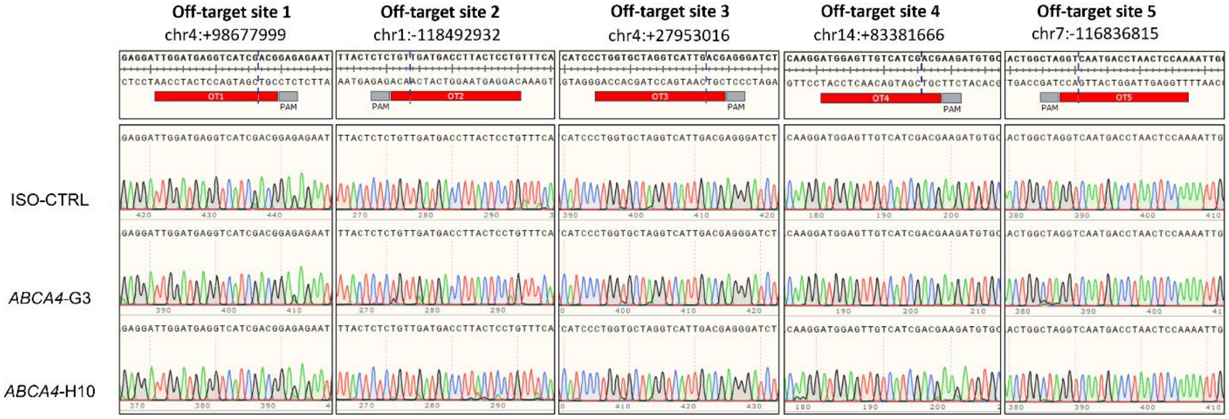


**Figure S2: Off-target mutagenesis analysis of *ABCA4* hiPSCs subclones.** (A) Sanger sequencing of the genomic regions containing the top 5 off-target sites associated to the gRNA used in this study as predicted by the IDT "CRISPR-Cas9 guide RNA design checker" tool. Chromatograms indicate no indels around the expected cut site (dashed line in blue).

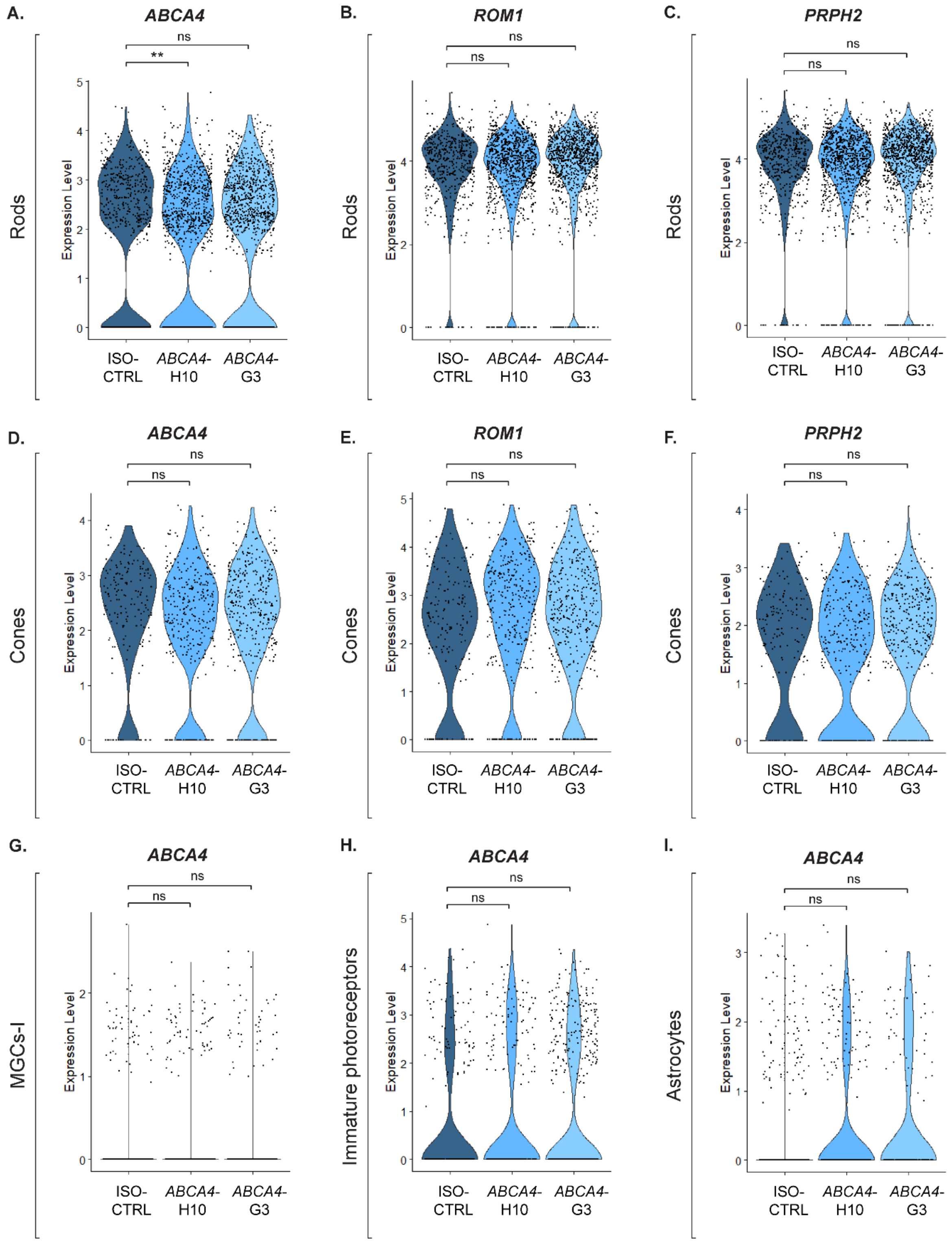


**Figure S3: Expression of *ABCA4*, *ROM1* and *PRPH2* in selected clusters of *ABCA4*-mutant and isogenic control organoids.** (A-C) Violin plots comparing the expression levels of *ABCA4* (A), *ROM1* (B), and *PRPH2* (C) in control and mutant rod photoreceptor cells. Significance was indicated as $p < 0.01$ (**), and ns (not significant). Adjusted p-values = 0,004 (**); all others = 1. (D-F) Violin plots comparing the expression levels of *ABCA4* (D), *ROM1* (E), and *PRPH2* (F) in control and mutant cone photoreceptor cells (all adjusted p-values = 1). (G-I) Violin plots comparing the expression levels of *ABCA4* in control and mutant Müller glial cells I (G), immature photoreceptors (H), and astrocytes (I) (all adjusted p-values = 1). Number of organoids used: ISO-CTRL *n* = 4; *ABCA4*-H10 *n* = 5; *ABCA4*-G3 *n* = 5 from 14 scRNA-Seq of the same differentiation round. Differential gene expression was evaluated at the single-cell level using Wilcoxon rank-sum test, with multiple-testing correction performed using the Benjamini–Hochberg method. Genes with an adjusted p-value < 0.05 were considered significantly differentially expressed.

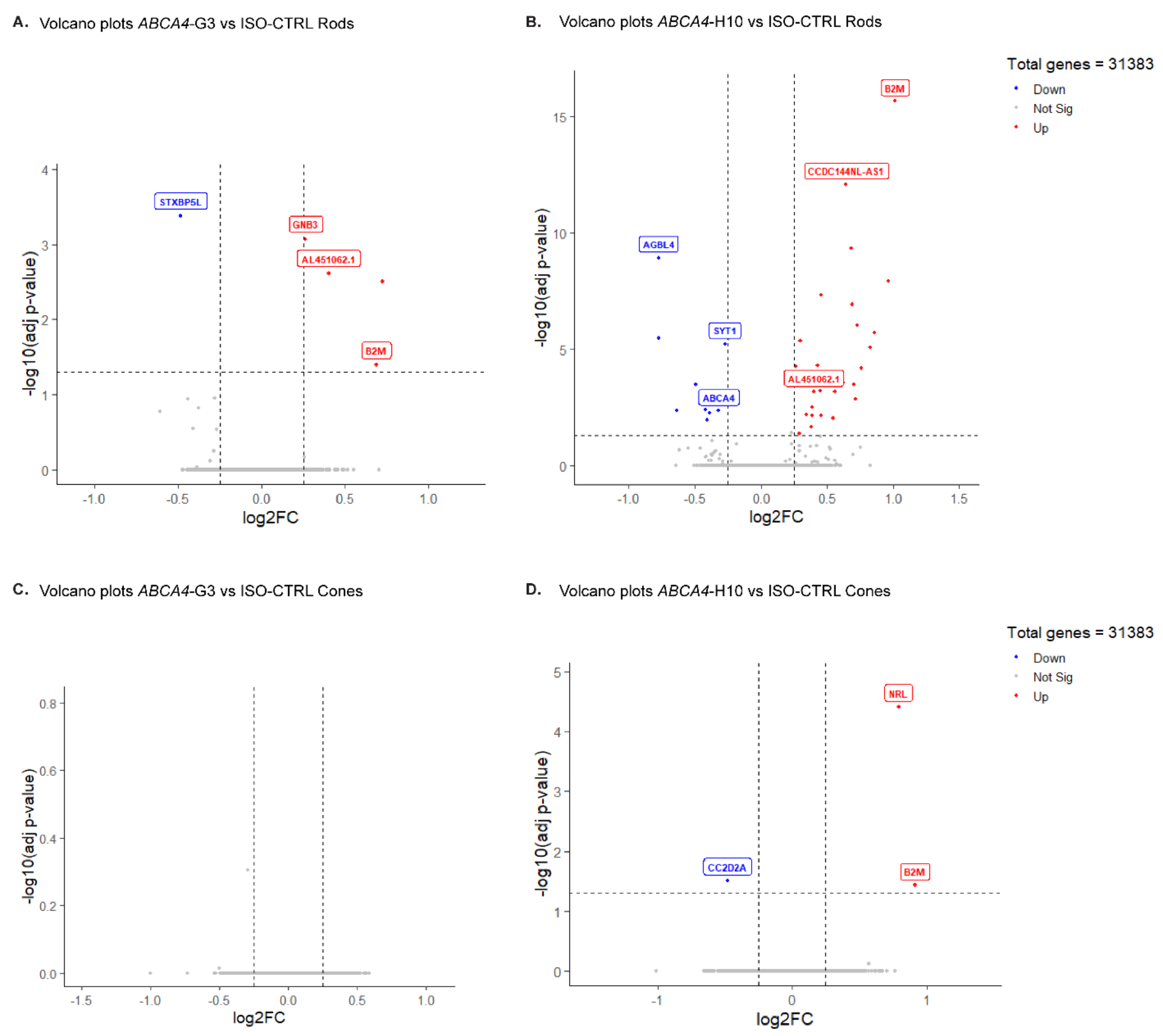


**Figure S4: Volcano plots showing differentially expressed genes in rods and cones of *ABCA4* subclones.** (A,B) Volcano plots indicating the downregulated (blue) and upregulated (red) genes in the rod photoreceptors of *ABCA4*-G3 (A) and *ABCA4*-H10 (B) compared to ISO-CTRL. (C,D) Volcano plots indicating the downregulated (blue) and upregulated (red) genes in the cone photoreceptors of *ABCA4*-G3 (C) and *ABCA4*-H10 (D) compared to ISO-CTRL. Statistical parameters: Wilcoxon rank-sum test, adjusted $p$ value $< 0.05$, absolute log2FC threshold = 0.25. Number of organoids used: ISO-CTRL $n = 4$; *ABCA4*-H10 $n = 5$; *ABCA4*-G3 $n = 5$ from 14 scRNA-Seq of the same differentiation round.

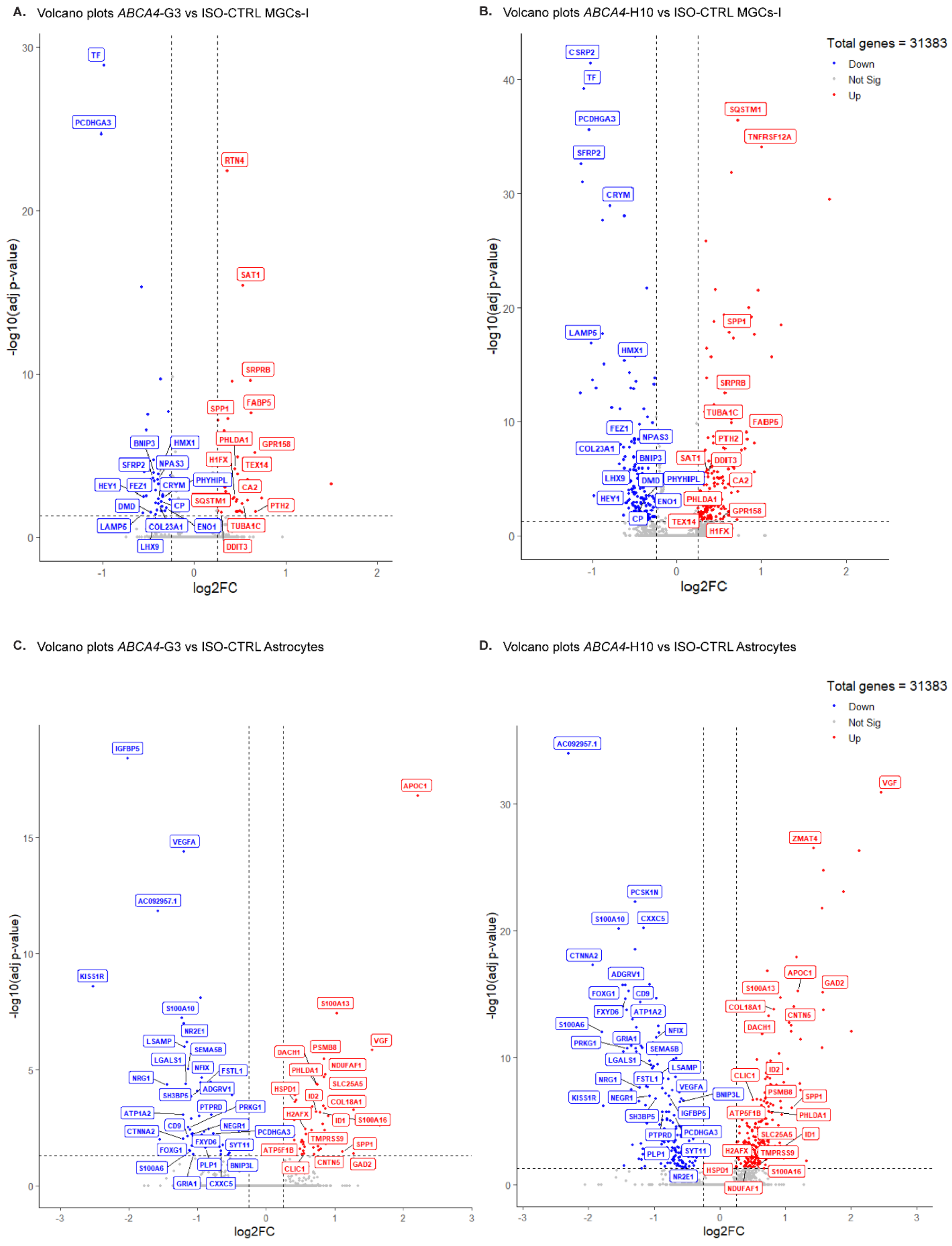


**Figure S5: Volcano plots showing differentially expressed genes in MGCs-I and astrocytes of *ABCA4* subclones.** (A,B) Volcano plots indicating the downregulated (blue) and upregulated (red) genes in the MGCs-I of *ABCA4*-G3 (A)

and *ABCA4*-H10 (B) compared to ISO-CTRL. (C,D) Volcano plots indicating the downregulated (blue) and upregulated (red) genes in the astrocytes of *ABCA4*-G3 (C) and *ABCA4*-H10 (D) compared to ISO-CTRL. Statistical parameters: Wilcoxon rank-sum test, adjusted $p$ value < 0.05, absolute log2FC threshold = 0.25. Number of organoids used: ISO-CTRL $n$ = 4; *ABCA4*-H10 $n$ = 5; *ABCA4*-G3 $n$ = 5 from 14 scRNA-Seq of the same differentiation round. Genes with an adjusted p-value < 0.05 were considered significantly differentially expressed.

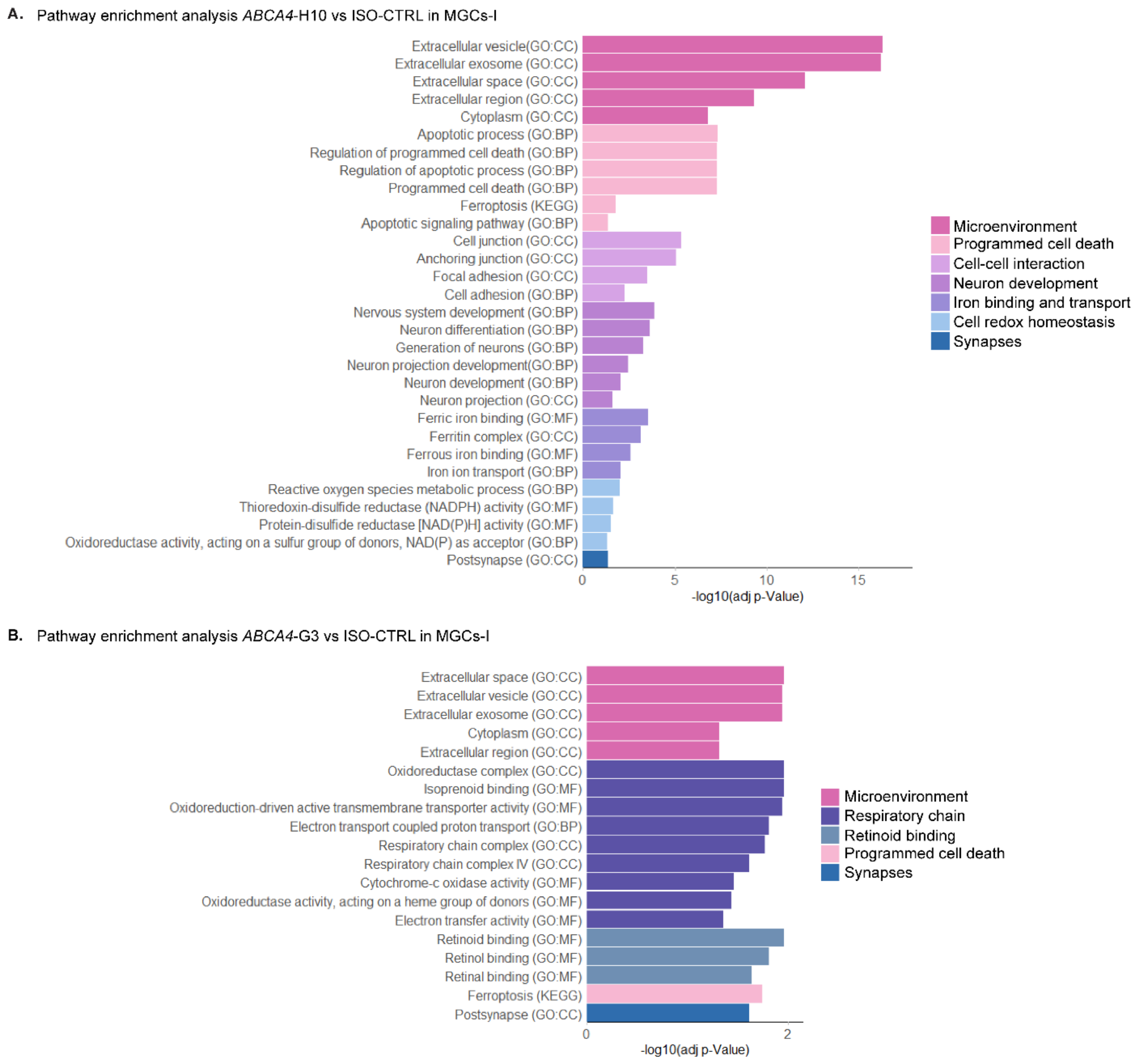


**Figure S6: Pathway enrichment analysis of *ABCA4* MGCs-I.** (A,B) Bar plots showing the most significant pathway enrichment terms found in the MGCs-I cluster of *ABCA4*-H10 (A) and *ABCA4*-G3 (B). Similar terms have been grouped into categories labeled with colors. Number of organoids used: ISO-CTRL *n* = 4; *ABCA4*-H10 *n* = 5; *ABCA4*-G3 *n* = 5 from 14 scRNA-Seq of the same differentiation round.

DEGs related to the pathway enrichment terms in the *ABCA4*-mutant subclones vs ISO-CTRL in MGCs-I

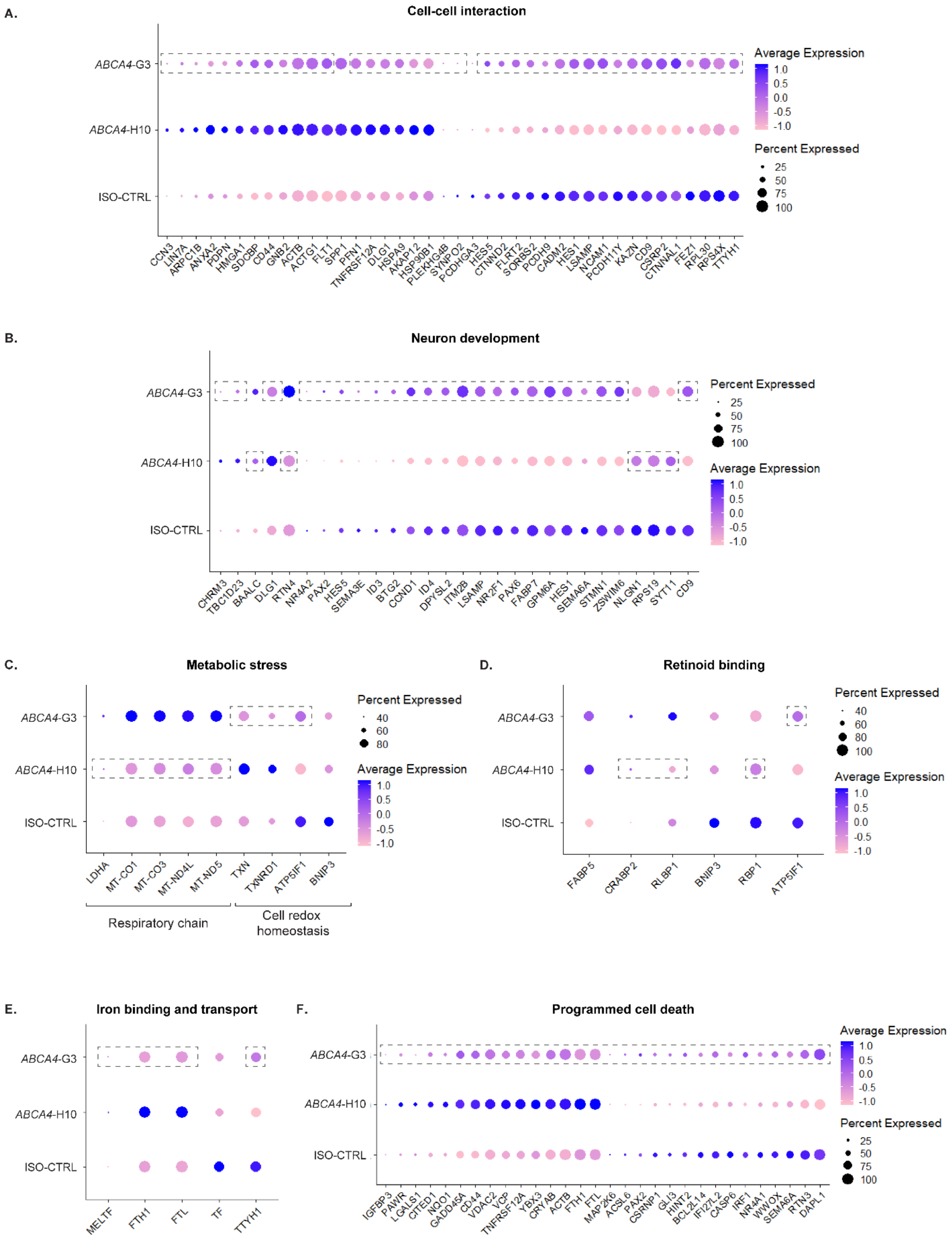

**Figure S7: Differential expression analysis of *ABCA4* mutant MGCs-I compared to isogenic controls.** Dot plots displaying DEGs associated with cell-cell interaction (A), neuron development (B), metabolic stress (C), retinoid binding (D), iron binding and transport (E), and programmed cell death (F). Dot plots (B) and (D) display additional DEGs retrieved from neuron development and programmed cell death terms that were not included in Figure 5, which only shows DEGs shared between subclones. The genes shown here represent subclone-specific DEGs. Dashed boxes highlight genes that did not exhibit differential expression in a specific subclone compared to ISO-CTRL. Number of organoids used: ISO-CTRL *n* = 4; *ABCA4*-H10 *n* = 5; *ABCA4*-G3 *n* = 5 from 14 scRNA-Seq of the same differentiation round. Differential gene expression was evaluated at the single-cell level using Wilcoxon rank-sum test, with multiple-testing correction performed using the Benjamini–Hochberg method. Genes with an adjusted p-value < 0.05 were considered significantly differentially expressed.

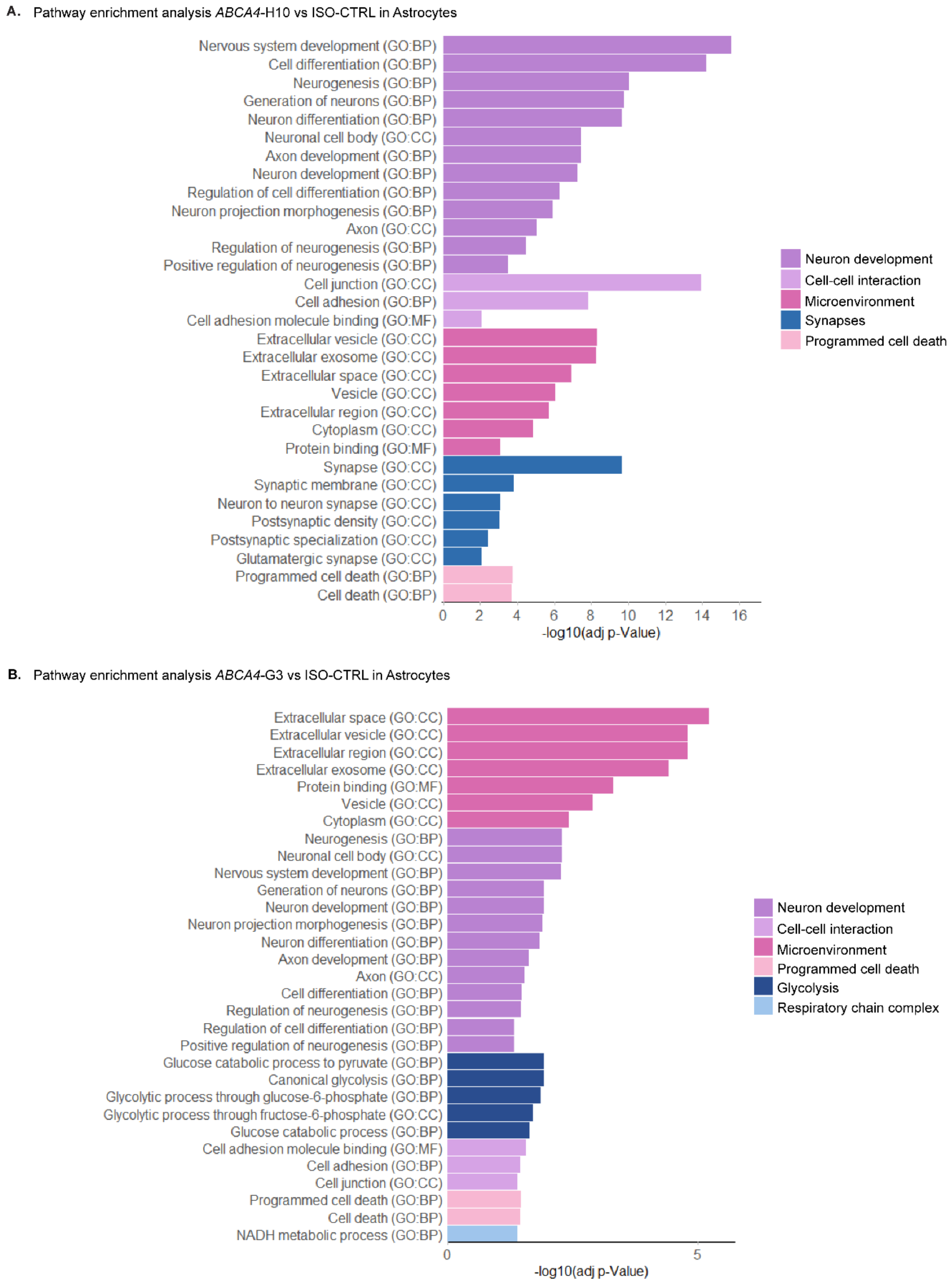


**Figure S8: Pathway enrichment analysis of *ABCA4* astrocytes.** (A,B) Bar plots showing the most significant pathway enrichment terms found in the astrocytes of *ABCA4*-H10 (A) and *ABCA4*-G3 (B). Similar terms have been grouped into categories labeled with colors. Number of organoids used: ISO-CTRL *n* = 4; *ABCA4*-H10 *n* = 5; *ABCA4*-G3 *n* = 5 from 14 scRNA-Seq of the same differentiation round.

DEGs related to the pathway enrichment terms in the *ABCA4*-mutant subclones vs ISO-CTRL in Astrocytes

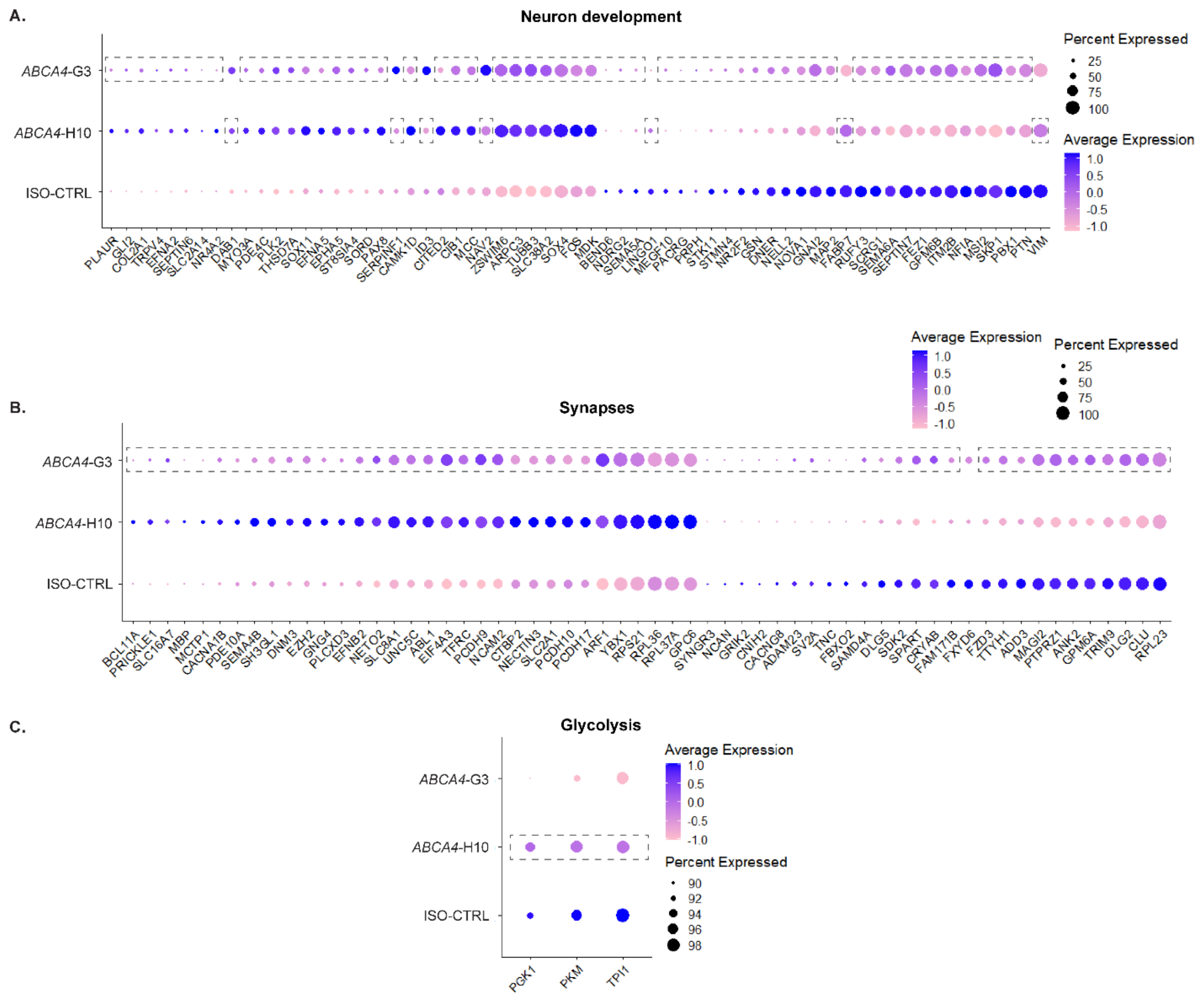


**Figure S9: Differential expression analysis of *ABCA4* mutant astrocytes compared to isogenic controls.** Dot plots displaying DEGs associated with neuron development (A), synapses (B), and glycolysis (C) that were not included in Figure 7, which only shows DEGs shared between subclones. The genes shown here represent subclone-specific DEGs. Dashed boxes highlight genes that did not exhibit differential expression in a specific subclone compared to ISO-CTRL. Number of organoids used: ISO-CTRL *n* = 4; *ABCA4*-H10 *n* = 5; *ABCA4*-G3 *n* = 5 from 14 scRNA-Seq of the same differentiation round. Differential gene expression was evaluated at the single-cell level using Wilcoxon rank-sum test, with multiple-testing correction performed using the Benjamini–Hochberg method. Genes with an adjusted p-value < 0.05 were considered significantly differentially expressed.

**Table S1:** hiPSC line information.

| Line name | Description | Gender |
|---|---|---|
| **LUMC0004iCTRL10 = ISO-CTRL**<br>**(hPSCreg name: LUMCi029-B)** | Control parental hiPSC line | Male |
| **LUMC0004iCTRL10_*ABCA4*$^{24KO}$ CLG3 = *ABCA4*-G3** | A 10-bp deletion–insertion introduces a premature stop codon in exon-24 of *ABCA4*.<br>c.3677-3686delinsTGA; p.(Val1192Ter) | Male |
| **LUMC0004iCTRL10_*ABCA4*$^{24KO}$ CLH10 = *ABCA4*-H10** | A 10-bp deletion–insertion introduces a premature stop codon in exon-24 of *ABCA4*.<br>c.3677-3686delinsTGA; p.(Val1192Ter) | Male |

Information on the hiPSC lines used in this study.

**Table S2:** Information related to the crRNA and donor template used to generate the *ABCA4*-mutant hiPSC lines and primers to assess the correct genomic mutation and absence of off-target mutagenesis.

| Name | Sequence (5'-3') |
|---|---|
| ***ABCA4* exon-24 crRNA** | CTGGAGTTAGGTCATCGACG |
| ***ABCA4* ssODN** | AGTCATCCCATCCATCTGTTGCAGGGGACCTGCAGCTGCTC GTCTAAGGGTTTCTCCACCACGTGTCCAGCCCACTGATAAC TCCAGAACAAGTCCTGGATGGTAAGGACTGGACGGGCCA TACTTGGGTTCCGTCTGGCAGCCATCTCCCAG |
| ***ABCA4* exon-24 Fw primer** | GACCCTTGGAGGTTCTGCTC |
| ***ABCA4* exon-24 Rv primer** | TGCATCACCATGCTTCCACT |
| **Off-target 1 (chr4:+98677999) Fw primer** | AGGGTACAGGTGAGGGGAAA |
| **Off-target 1 (chr4:+98677999) Rv primer** | TCTTCATGTTGGTGAGCCCC |
| **Off-target 2 (chr1:-118492932) Fw primer** | ACTCCCTTTGTCAGCTCTACA |
| **Off-target 2 (chr1:-118492932) Rv primer** | ACACTAGGTATTGTTTAGTTGGTGT |
| **Off-target 3 (chr4:+27953016) Fw primer** | CAGCAGGTAACAGGCACTCA |
| **Off-target 3 (chr4:+27953016) Rv primer** | TTTCTTCATAGGTGAACAACTGTC |
| **Off-target 4 (chr14:+83381666) Fw primer** | CCATTGGAGGGAAAGATACACTAAG |
| **Off-target 4 (chr14:+83381666) Rv primer** | TCCAAGTGATTTTGTCTAGTATCATGA |
| **Off-target 5 (chr7:-116836815) Fw primer** | CCTGTTGGGAGAATGCTGGT |
| **Off-target 5 (chr7:-116836815) Rv primer** | TTTGATCCCCGAACCACACC |

Forward (Fw) and Reverse (Rv) primers used to confirm the CRISPR/Cas9-mediated insertion of the premature stop codon in *ABCA4* exon-24 and the absence of mutations in the top 5 predicted sites.

**Table S3:** List of materials used in this study.

| Materials | Source | Identifier |
|---|---|---|
| **tracrRNA** | IDT | 1072532 |
| **Nuclease-Free Duplex Buffer** | IDT | 11-01-03-01 |
| **SpCas9 Nuclease V3** | IDT | 1081058 |
| **Neon™ Transfection System kit** | Invitrogen | MPK10096 |
| **Neon™ Transfection System** | Invitrogen | MPK1025 |
| **Matrigel hESC-Qualified Matrix** | Corning | 354277 |
| **mTeSR plus medium** | STEMCELL Technologies | 100-0276 |
| **Accumax** | STEMCELL Technologies | 07921 |
| **Gentle Cell Dissociation reagent** | STEMCELL Technologies | 100-0485 |
| **CloneR** | STEMCELL Technologies | 05888 |
| **Cryostor** | STEMCELL Technologies | 07930 |
| **DMEM/F12** | Life Technologies | 11320074 |
| **DMEM(1X) + GlutaMAX** | ThermoScientific | 10569010 |
| **Fasudil HCl** | Focus Biomolecules | 10-2137 |
| **40 µm cell strainer** | pluriSelect | 43-10040-40 |
| **Blebbistatin** | abcam | ab120425 |
| **Micro-molds** | Merck | Z764000-6EA |
| **MEM NEAA 100X** | Life technologies | 11140-035 |
| **Taurine** | Merck | T0625 |
| **Neurocult SM1 50X** | STEMCELL Technologies | 05711 |
| **N2 supplement 100X** | Life technologies | 17502048 |
| **Heparin** | Merck | H-9399 |
| **Smoothened agonist (SAG)** | Selleck Chemicals | S7779 |

| | | |
|---|---|---|
| **Gamma secretase inhibitor IX (DAPT)** | Selleck Chemicals | S2215 |
| **Fetal Bovine Serum** | Serana | S-FBS-CO-015 |
| **Poloxamer 188** | Merck | P5556 |
| **Retinoic acid** | Merck | R-2625 |
| **Antibiotic-antimycotic 100X** | Merck | A5955 |
| **Tissue-Tek O.C.T. Compound** | Sakura Finetek | 4583 |
| **Vectashield Antifade Mounting Medium** | Vector Laboratories | H-1800-10 |
| **Papain Dissociation kit** | Worthington | I-LK 03150 |

**Table S4:** List of antibodies used in this study.

| Antibody | Dilution | Source | Identifier |
|---|---|---|---|
| **Anti-CRALBP** | 1:500 | Abcam | ab15051 |
| **Anti-Recoverin** | 1:600 | Millipore | AB5585 |
| **Anti-MUPP1** | 1:150 | BD Transduction Laboratories | M98820 |
| **Anti-Rhodopsin** | 1:500 | Millipore | MAB5356 |
| **Anti-R/G Opsin** | 1:300 | Millipore | AB5405 |
| **Anti-INPP5E** | 1:200 | Proteintech | 17797-1-AP |
| **Anti-ABCA4 (N-terminal)** | 1:1000 | Millipore | MABN2440 |
| **Anti-ROM1** | 1:300 | Proteintech | 21984-1-AP |
| **Anti-PRPH2** | 1:400 | Proteintech | 18109-1-AP |
| **Goat anti-rabbit IgG (H+L) Highly Cross-Adsorbed Secondary Antibody, Alexa Fluor 488** | 1:1000 | Invitrogen | A-11034 |
| **Goat anti-mouse IgG H&L, Alexa Fluor 555** | 1:1000 | Abcam | ab150118 |

Product information about the antibodies used in the immunohistochemical analyses.

**Supplementary Methods**

***1. Karyotyping analysis***

Normal karyotypes were confirmed by G-banding with 20 metaphases analyzed per cell line. Analysis was performed by the Section Genome Diagnostics, Clinical Genetics, LUMC, using the Leica Biosystems CytoVision (Leica Microsystems, Wetzlar, Germany).

***2. Copy number variation analysis***

The genomic DNA of hiPSC lines was analyzed for copy number variations using the iCS-digital™ PSC 24-probes kit (Stemgenomics, Montpellier, France).

***3. Short Tandem Repeat (STR) analysis***

STR-analysis was performed by the LUMC Forensic Laboratory for DNA-research (FLDO), using the PowerPlex Fusion System 5C autosomal STR kit (Promega Corporation, Madison, WI, USA).

***4. Transmission electron microscopy (TEM) analysis***

Samples were fixed at room temperature in 1.5% glutaraldehyde prepared in 0.1 M cacodylate buffer by adding an equal volume of double-strength fixative directly to the culture medium for 1 hour. Following fixation, samples were washed three times in 0.1 M cacodylate buffer and post-fixed for 1 hour on ice in 1% osmium tetroxide supplemented with 1.5% potassium ferricyanide in the same buffer. After three additional buffer washes, samples were dehydrated through graded ethanol solutions, followed by acetone and increasing concentrations of EPON resin (LX112), and finally infiltrated with 100% EPON. Organoids were embedded in molds filled with EPON and polymerized at 70 °C for 48 hours. Ultrathin sections (90 nm) were cut using a Reichert Ultracut S ultramicrotome (Leica Microsystems, Wetzlar, Germany), contrasted with uranyl acetate and lead citrate, and imaged using a Tecnai T12 transmission electron microscope operated at 120 kV and equipped with a OneView camera (ThermoFisher Scientific, Waltham, MA). Serial images were acquired and stitched into composite micrographs and analyzed on ImageScope (v12.4.6).

***5. scRNA-Seq computational analysis***

Single cells were assigned to their sample of origin based on hashtag oligonucleotide (HTO) enrichment. Cells positive for a single HTO were classified as singlets and retained for further analysis. Quality control filtering was applied to keep only cells with nFeature_RNA between 500 and 8000, nCount_ RNA < 40000, and percent.mt <15. Data were normalized using the LogNormalize method with a scale factor of 30000, and the 2000 most variable genes were identified. Principal component analysis (PCA) was performed on scaled expression values, and the first 15 principal components were used for clustering. Nineteen transcriptionally distinct clusters were identified and visualized using UMAP. Cluster identities were assigned based on established retinal cell type marker genes identified using the FindAllMarkers function in Seurat. For downstream analyses, data were subset by cluster and experimental condition, and differential gene expression was assessed using Wilcoxon rank-sum test. Genes with an adjusted p-value below 0.05 were subjected to pathway enrichment analysis using gProfiler. In dot plots, dot size represents the proportion of cells expressing a given gene within each cluster, while color intensity reflects the average expression level.
